# Natural prey capture reveals flexible reconfiguration of canonical motor-cortical dynamics for online control

**DOI:** 10.64898/2026.09.01.748436

**Authors:** Jeffrey D. Walker, Paul Aparicio, Shreya Saxena, Nicholas G. Hatsopoulos, Jason N. MacLean

**Author notes:** Co-senior authors (Nicholas G. Hatsopoulos, Jason N. MacLean).

## Abstract

The ability to record neural populations during natural behavior now allows us to ask whether canonical motor-cortical dynamics, defined largely in constrained reaches to static targets, also organize self-paced, feedback-rich actions that require continuous correction. We recorded sensorimotor cortex in marmosets freely capturing live prey, a behavior that separates into ballistic quick captures and dynamic pursuits requiring real-time adjustment. Ballistic captures corresponded to canonical cortical reaching dynamics without an instructed delay, including near-orthogonal premovement and movement subspaces. Pursuits did not recruit additional neural dimensions; despite their elaborate kinematics, dynamics evolved within the same low-dimensional subspace. Corrective movements during pursuit reorganized this geometry into a distinct feedback-dominated regime, in which premovement activity weakly predicted movement, trajectory tangling increased, and a smaller, slower rotation restarted from a shifted initial state. Yet all movement types shared a sequential firing-rate backbone, which during ballistic capture was organized across the cortex as a somatotopically directed propagating wave that weakened when movements required online correction. These results show that sensorimotor cortex preserves canonical dynamical structure while flexibly reconfiguring it for online control in natural behavior.

## Introduction

To capture moving prey, a predator must prepare an action, launch it, and revise it in real time as the target performs evasive maneuvers. This closed-loop control problem is central to natural movement, yet it is largely absent from the constrained reaching tasks that have shaped current models of motor cortical dynamics. The common marmoset (*Callithrix jacchus*) provides the opportunity to study this problem directly: it captures prey voluntarily, without laboratory training (Walker et al. 2020), while using reach to grasp movements that are refined over development as juveniles learn to hunt (Schiel et al. 2010), and hunting occupies a substantial fraction of the marmoset’s daily activity (Rylands and de Faria 1993; Stevenson and Rylands 1988). Marmosets also adapt their strategy to the prey itself: fast, evasive insects evoke visually guided reach-to-grasp movements, whereas slower ground-dwelling prey are taken directly by mouth (Ngo et al. 2022). A single pursuit can thus combine premovement preparation, ballistic execution, and rapid online correction with prediction as the target evades (Hein et al. 2020; Shaw et al. 2023), making prey capture a natural arena for testing whether canonical motor-cortical dynamics generalize to the flexible control demands of real-world action.

How sensorimotor cortex produces this flexibility remains unclear because most accounts of cortical control of the primate forelimb come from a very different behavioral regime. Standard paradigms rely on stereotyped reaches to static or abruptly displaced targets, externally paced and repeated across many trials. By design these tasks isolate movement from the continuous sensory feedback that structure natural action. This approach has been highly productive, revealing principles from single-neuron directional and trajectory tuning (Georgopoulos et al. 1986; Archambault et al. 2011; Hatsopoulos et al. 2007) to low-dimensional population dynamics that organize movement preparation and execution (Kaufman et al. 2014; Elsayed et al. 2016; Saxena and Cunningham 2019). Yet these signatures were established largely under constrained, repetitive conditions. It remains unknown whether they reflect general principles of sensorimotor cortical control or instead the task structure that revealed them. This distinction is critical for understanding whether motor-cortical dynamics generalize beyond the laboratory to the self-paced, feedback-driven demands of natural behavior.

Motor cortex is organized spatially as well as dynamically. Beyond its low-dimensional population geometry, the forelimb representation is laid out across the cortical sheet as a somatotopic gradient, and movement-related activity can travel across this sheet as a propagating cortical wave (Rubino et al. 2006; Balasubramanian et al. 2020; Liang et al. 2023). Whether such propagation follows the somatotopic map rather than a fixed anatomical axis, as a proximal-to-distal organization would require, has remained unresolved. How this spatial organization participates in natural, feedback-driven action, and whether it tracks the same shifts in operating regime that appear in the population dynamics, is unknown.

To test this directly, we employed a prey-capture paradigm in which freely moving marmosets voluntarily pursue and capture live moths executing unpredictable, evasive maneuvers, while we recorded wirelessly from sensorimotor cortex. This behavior spans a natural continuum of forelimb control, from rapid captures completed in a single ballistic reach to extended pursuits in which abrupt prey evasions demand multiple sequential corrective movements. We used this natural behavioral variability to ask whether the canonical population-dynamical signatures of primate motor cortex persist during feedback-rich natural behavior, how these dynamics are altered when action must be corrected online, and how the underlying recruitment is organized across the cortical sheet. We find that prey capture preserves a shared low-dimensional dynamical structure across movement types, but that corrective movements shift sensorimotor cortex into a distinct operating regime, a shift mirrored in the coherence of a somatotopically directed cortical wave. Thus, natural behavior reveals canonical motor cortical dynamics as flexible structures, reconfigurable for continuous control rather than fixed templates for movement.

## Results

### Natural prey capture evokes ballistic reaches and feedback-driven corrective movements

To examine sensorimotor cortical dynamics during natural behavior, we developed a voluntary prey-capture task based on the marmoset’s natural behavioral repertoire (Fig. 1A). When an animal adopted a poised posture at the edge of the workspace, a live moth was released from a port and typically landed on the workspace floor. Capture attempts began after the moth landed and followed a relatively stereotyped initial kinematic sequence: hand lift, extension, grasp, and retraction. Some attempts were completed on the first reach, before the moth executed an evasive maneuver; we refer to these episodes as quick captures (Fig. 1B,D). Failed initial attempts instead led to extended pursuits, during which marmosets either made sequential corrective movements with the arm at extension or fully retracted and reset the hand before launching another reach. (Fig. 1C,E). We recorded behavior with high-speed cameras, reconstructed three-dimensional hand position using markerless keypoint tracking, and used an algorithmic segmentation procedure to identify movement events of each type (see Methods). In total we analyzed 72 prey-capture episodes in marmoset TY (comprising 217 segmented movement events) and 35 episodes in marmoset JL (124 events).

**Fig. 1.**
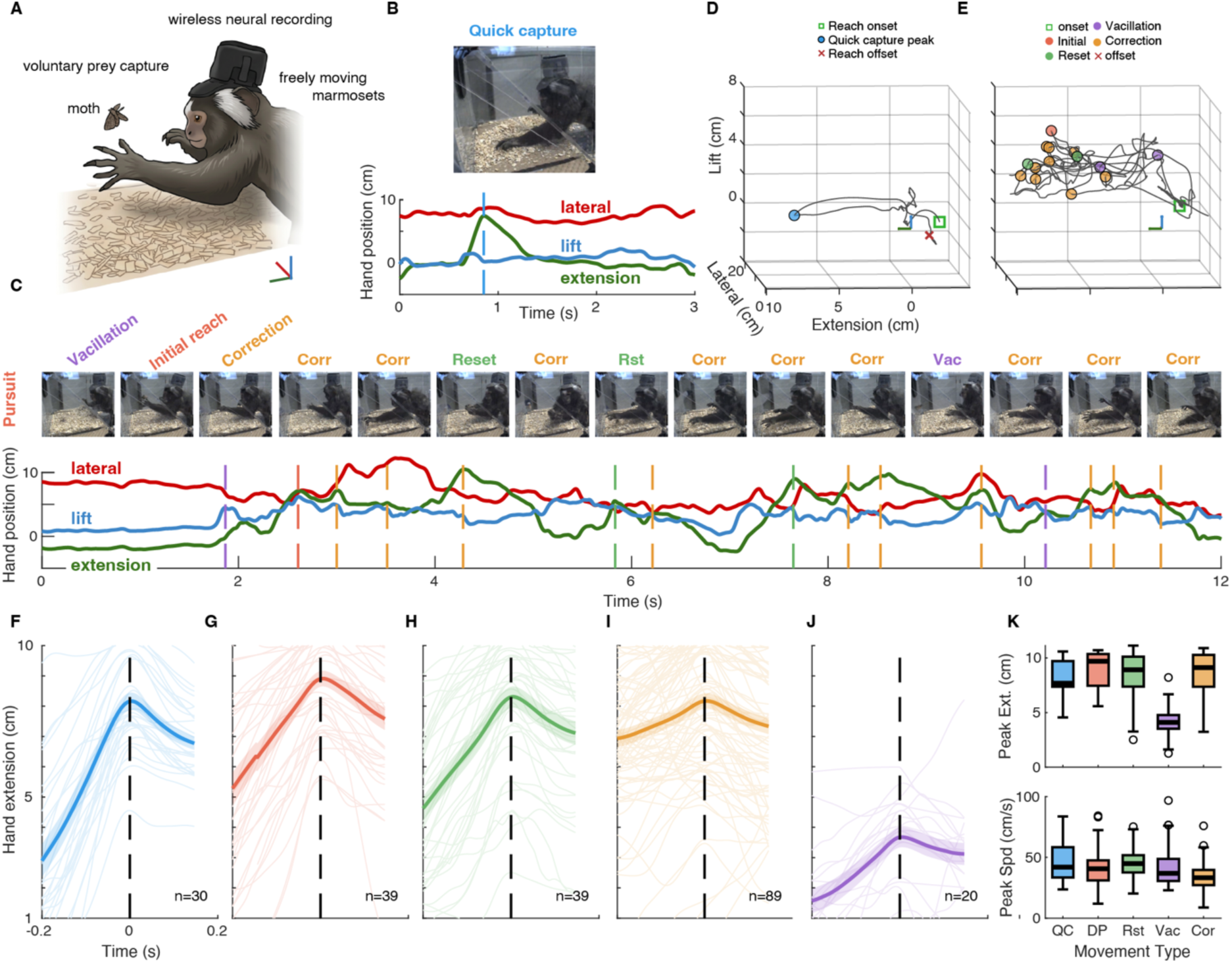
Marmoset prey capture behavior: quick captures and dynamic pursuits. A) Illustration of prey capture experimental setup, with freely moving marmosets engaging in voluntary prey capture and concurrent wireless neural population recordings from sensorimotor cortex. B) Hand kinematics from a quick capture episode where moth is captured with single ballistic reach with video frame at peak hand extension. C) Hand kinematics from an episode of pursuit where the marmoset expresses multiple sequential corrective movements and reach resets before capturing the moth. Filmstrip illustrating frames corresponding to each movement type: initial reach, reach resets, corrections and vacillations. D) 3D hand position in prey capture workspace for example quick capture episode in B. E) Hand kinematics from example episode of pursuit in C. F-J) Mean hand extension across movement types with individual event traces in thinner lighter lines for (F) quick capture, G) initial dynamic pursuit reach, H) reach resets, I) corrections and J) vacillations. K) Distributions of peak extension and peak hand speed across movement types. Init: initial reach. Vac: vacillation. Corr: correction or corrective movement. Rst: reach reset or reset.

We next classified prey-capture movements according to their kinematic structure. Initial reaches during quick captures closely resembled the first reaches of pursuit episodes and the subsequent reach resets made during pursuit (Fig. 1F-H). We therefore parsed prey capture into movement classes: quick capture reaches (Fig. 1F), the first reaches of pursuit episodes (Fig. 1G), resets (Fig. 1H), in which the marmoset retracted the hand before initiating another reach, online corrections (Fig. 1I) executed while the hand was already extended, and vacillations (Fig. 1J), incomplete reaches in which the hand did not reach full extension. Peak hand extension was comparable across movement types with the exception of vacillations (Fig. 1J; 1K). Corrections were markedly smaller and slower than the initial reaches that they followed, as expected from movements initiated with the hand already extended: median correction reach extent ≈ 3.0 cm vs ≈ 7.3 cm in TY; ≈ 2.7 vs 7.6 cm in JL. Thus, any increased neural variability associated with corrections cannot be explained by corrections being larger or faster movements. Although initial reaches were kinematically similar across quick captures and pursuits, pursuit episodes were more variable and occupied a larger portion of the workspace. Kinematic variance at the firsthand extension peak was consistently greater during pursuits than during quick captures (Fig. 1D,E) and lateral hand position variance increased across successive extension peaks within episodes of dynamic pursuit (Fig. 1D,E). Thus, prey capture produced both stereotyped ballistic reaches and more variable corrective sequences within a single natural behavior. Despite this variability in episode structure (Fig. 1B,C), hand kinematics provided a robust behavioral substrate for linking sensorimotor cortical population activity to natural prey capture.

### Quick captures express canonical motor-cortical dynamics

Given the natural separation of prey capture into quick captures and extended pursuits, we first examined sensorimotor cortical neural population dynamics during quick captures. This allowed us to test whether a natural self-initiated capture movement expresses canonical motor cortical dynamics previously established in highly constrained reaching tasks including instructed-delay reaches and center-out movements.

In our task, the marmoset typically reached for the moth immediately after the moth landed on the workspace floor. During quick captures, neural population activity followed structured trajectories that aligned with behavioral sequences (Fig. 2A). At the start of each episode, the neural state remained near a consistent origin point, then shifted after the moth landed, before the onset of hand movement. The largest transition occurred with hand extension and was accompanied by sequential population activity across much of the recorded population. When units within each area were ordered by their latency to peak response during quick captures, this sequence revealed the major behavioral phases of the capture episode (Fig. 2A). In a low dimensional population space identified by PCA of soft normalized firing rates (see Methods) the same structure appeared as a broad trajectory through state space reaching an apex coincident near peak hand extension (Fig. 2A,B). During hand retraction the neural state returned toward the origin point while also expressing trajectory features corresponding to hand to mouth movements and prey manipulation around the mouth (Fig. 2A,B). Clustering of behavioral and neural states further revealed consistent structure in both physical space and neural space across episodes and marmosets (Fig. 2B).

**Fig. 2.**
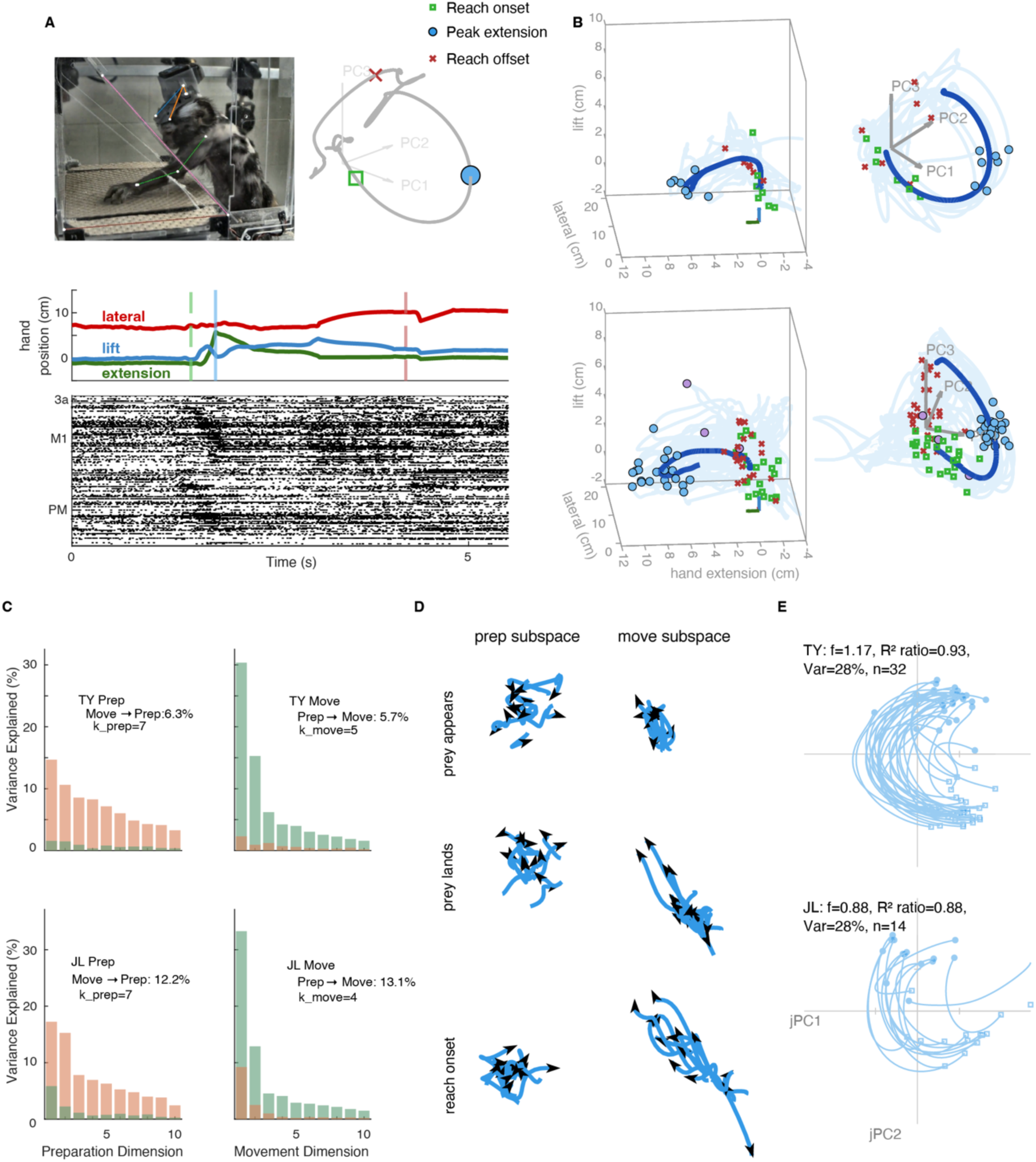
Sensorimotor cortical population dynamics during quick capture episodes. A) Data from example quick capture episode data. Top left: Frame of video at peak extension. Middle: 3D hand kinematics with vertical lines correspond to reach onset, peak extension and reach offset. Bottom: Raster of sensorimotor cortical unit spiking with units ordered according to recruitment latency. Top right: Behavioral state annotated neural trajectory. B) All quick capture hand (left) and neural (right) trajectories for both marmosets (top: JL, bottom: TY). C) Percentage of variance explained by preparatory and movement subspaces during quick captures illustrate that preparatory and movement activity is near orthogonal during quick captures. D) Activation within the first two dimensions of the preparatory (top) and movement (bottom) subspaces when prey appears, when prey lands and at the onset of marmoset’s reach for TY. E) Phase portraits of neural trajectories for quick capture episodes for both marmosets in the plane of jPC1 and jPC2 illustrating rotational dynamics during relatively simple reach to grasp.

We next asked whether these natural capture population dynamics contained canonical signatures of motor cortical dynamics: near orthogonal preparatory and movement-related activity, and rotational population dynamics. To quantify preparatory and movement subspaces, we adapted the cross-subspace variance approach introduced by Elsayed and colleagues (Elsayed et al. 2016, 2017). For quick captures we defined the preparatory epoch as 500 ms to 200 ms prior to movement onset, and the movement epoch as 100 ms prior to movement onset to peak hand extension. Even though the task contained no instructed delay and the animals reached as soon as the prey was within range, activity before and during movement occupied largely distinct subspaces. Cross-epoch variance explained was low in both marmosets: (TY cross variance: Prep→Move 5.7%, Move→Prep 6.3%; JL cross variance: Prep→Move 13.1%, Move→Prep 12.2%) (Fig. 2C). Thus, near-orthogonal preparatory and movement subspaces emerged during natural prey capture despite the absence of an experimentally imposed preparatory period, indicating that this canonical geometry is an intrinsic feature of motor-cortical control rather than a consequence of the instructed delays typically used to elicit it.

To visualize how these subspaces evolved during quick captures, we examined neural trajectories in the leading dimensions of the preparatory and movement subspaces, adapting the approach used by Lara and colleagues (Lara et al. 2018; Fig. 2D). Preparatory-subspace trajectories were broadly distributed when the prey appeared and became more consolidated by movement onset, whereas movement-subspace trajectories showed the opposite pattern, remaining clustered near the origin during prey appearance and landing before expanding sharply at movement onset (Fig. 2D).

Finally, we tested whether quick captures expressed rotational dynamics. Neural trajectories in principal component space exhibited a curved trajectory from movement onset to movement offset (Fig. 2A,B), which we quantified using jPCA (Churchland et al. 2012) in 400ms windows centered on kinematic events anchored to peak hand extension ([-0.25s, +0.15s]). Quick capture dynamics were well described by rotational structure with skew-symmetric dynamics accounting for most of the variance captured by the best-fit linear dynamics model (R² ratio = R²(M_skew)/R²(M_best) = 0.93 in TY and 0.88 in JL; Fig. 2E). The dominant rotation frequency was consistent across animals and matched the time scale of reaching behavior (1.17 Hz TY, 0.88 Hz JL). In both marmosets the first jPCA plane explained 28% of variance.

These analyses show that sensorimotor cortical dynamics during natural prey quick captures express canonical features of motor cortical population activity previously established in animals performing highly constrained reaching tasks. Even without an instructed delay, quick captures exhibited structured premovement activity, near-orthogonal preparatory and movement subspaces and strong rotational dynamics aligned with the unfolding reach. Thus, the canonical dynamical organization of motor cortex generalizes to a natural, self-initiated prey-capture behavior.

### Dynamic pursuit elaborates the quick-capture population geometry

Having established that quick captures express canonical reaching dynamics, we next asked how population dynamics are reorganized during the added complexity of dynamic pursuit. Specifically, we tested whether pursuit recruits new neural dimensions or instead reuses the quick-capture neural population geometry, and how online corrections are expressed within that shared geometry established above. Quick captures followed a stereotyped neural trajectory. Population activity began near a consistent starting region, shifted after the moth landed, and then underwent a large transition aligned with hand extension. This transition corresponded to sequential activation across the population and smooth rotational flow through the low dimensional state space. In episodes where the marmoset went on to inspect and eat the moth, the trajectory showed additional features corresponding to hand supination, hand-to-mouth movement, and return toward the initial state (Fig. 2A,B; Supplementary Video S1).

Dynamic pursuit episodes were behaviorally more complex. Hand trajectories occupied a larger portion of the workspace than quick captures and often contained multiple feedback-driven corrective movements (Fig. 1D,E; Fig. 3A,B). Nevertheless, neural trajectories during pursuit retained the broad rotational structure observed during quick captures, with additional trajectory features aligned to corrective movements. Extended reaches containing multiple corrective adjustments produced repeated smaller excursions through the low-dimensional subspace, with each excursion aligned to a discrete corrective movement (Fig. 3A,F; Supplementary video S1). Pursuit episodes that included reach resets, in which the marmoset retracted the hand toward the torso before launching another reach, produced larger sweeps through population space, often interleaved with smaller local rotations associated with fine adjustments during the extended phase of the reach (Fig. 3A,F; Supplementary video S1). These trajectories reveal how dynamic pursuit combines large state-space transitions associated with ballistic reach execution with smaller local dynamics associated with feedback-driven corrections. Initial ballistic reaches and subsequent corrective movements continued to engage the same units in a consistent temporal sequence. Thus, dynamic pursuit did not replace the quick-capture trajectory with a distinct population response but instead elaborated the same neural sequence through additional correction-related dynamics.

**Fig. 3.**
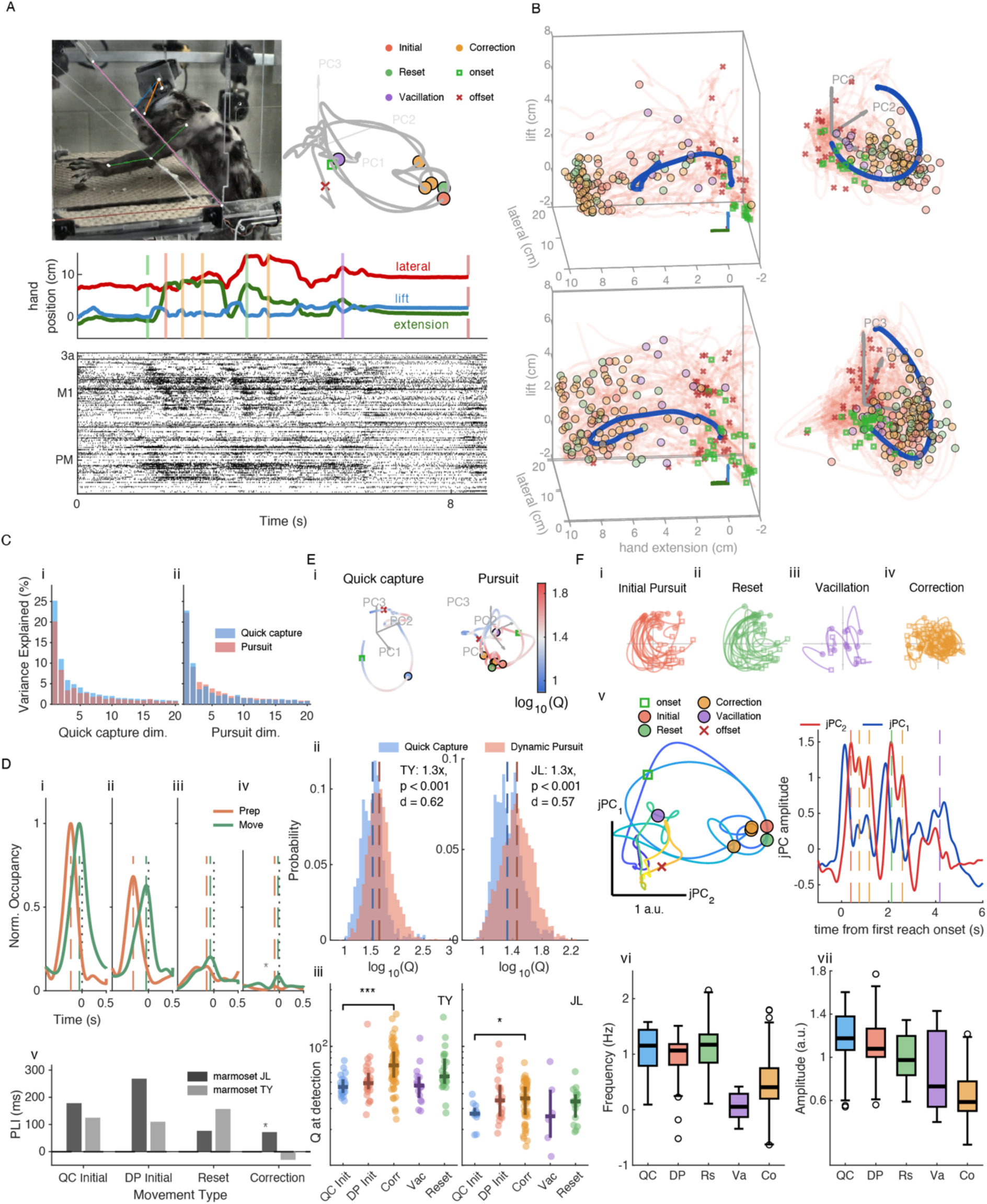
Dynamical features of neural population responses within sensorimotor cortex during complex episodes of prey capture pursuit. **A)** Example episode of dynamic pursuit with multiple corrective movements. Frame from video corresponding peak hand extension of the pursuit initiation. Hand kinematics (green axis is axis of extension, blue is axis of hand lifting, red is the hand moving either right or left). Raster of sensorimotor cortical population activity, ordered as in Fig. 2, with neural population dynamics for this episode. Note dynamical features corresponding to online corrections after initial sweep through space corresponding to initial ballistic reaching movement. The variability around the origin corresponds to pre and post movement epochs. **B)** 3D hand (left) and neural (right) trajectories for dynamic pursuit episodes both marmosets (top: JL, bottom: TY), only every other episode trajectory plotted for legibility. **C)** (i-ii) Cross-variance for subspaces derived from full quick capture and dynamic pursuit episodes for TY. **D)** (i-iv) Preparatory and movement subspace occupancy around movement type detections: (i) quick capture, (ii) pursuit initiation, (iii) reset and (iv) corrections. (v) Preparation lead index (PLI) across subjects and movement types. **E)** (i) Example quick capture and pursuit episode neural trajectories with tangling magnitude (log₁₀ Q) encoded in color. Colorscale centered between condition medians: JL [0.89, 1.89]. (ii) Distributions of log₁₀(Q) for quick-capture (blue) and dynamic-pursuit (red) episodes; both subjects. Dashed lines indicate medians. Two-sample KS test on log₁₀(Q): TY p < 10⁻¹⁰, KS D = 0.26, Cohen’s d = 0.62, n = 1,440 QC / 4,972 DP timepoints, DP/QC median ratio = 1.34×; JL p < 10⁻¹⁰, D = 0.27, d = 0.57, n = 2,295 / 4,850, ratio = 1.35×. (iii) Per-event tangling sampled from the episode-level Q cache at event detection time (window [−250, +150] ms; Q = per-event median). Bracket: QC-Initial vs Correction Wilcoxon rank-sum (TY p = 1.6 × 10⁻⁵, r = 0.46, median Q 45.6 vs 69.3; JL p = 2.0 × 10⁻², r = 0.29, median Q 27.1 vs 36.5). Event counts (TY / JL): QC-Init 20 / 12, DP-Init 30 / 21, Correction 67 / 52, Vacillation 16 / 6, Reset 24 / 18. **F)** (i-iv) Phase portraits within the first jPCA plane computed for each movement type (marmoset JL). (v) Example jPCA trajectory projected through the episode-scale basis, with jPC1/jPC2 time series, for the dynamic-pursuit episode in panel A. (vi) Per-event rotational frequency by movement type in the shared cross-type plane; box = median and IQR, whiskers 1.5×IQR, points = per-event values. Per-type point frequencies and 95% confidence intervals use the bootstrapped dominant M_skew eigenvalue (resampled within type, percentile intervals). (vii) Per-event detrended peak-to-peak rotational amplitude by movement type (quick capture, dynamic-pursuit initiation, reset, and correction in the shared plane; vacillation in its own plane, reported separately). Because events within a pursuit episode are not independent, significance is not marked on the panels; frequency (vi) and amplitude (vii) differences across movement types are evaluated with a linear mixed-effects model (movement type as a fixed effect; subject and episode as random effects) and an episode-block bootstrap as a non-parametric cross-check, reported in the text and Methods. The pre-specified quick-capture-versus-correction contrast is significant for both frequency (−0.506 Hz, p = 2×10⁻⁷; bootstrap 95% CI +0.35 to +0.69 Hz) and amplitude (−0.452 a.u., p = 1×10⁻⁹; bootstrap sign-concordant).

#### Dynamic pursuit reuses the quick-capture geometry but alters the timing of preparatory and movement dynamics

Given that pursuit behavior was more complex than quick-capture behavior, we first asked whether this added complexity reflected recruitment of new neural dimensions. We projected quick-capture and dynamic-pursuit population activity into subspaces derived from each condition and quantified cross-subspace variance explained. Dynamic pursuit responses were well explained by the quick-capture subspace and quick capture responses were likewise well explained by the pursuit derived subspace (Fig. 3C,i-ii). Thus, the more elaborate trajectories observed during pursuit did not reflect a wholesale expansion into new population dimensions. Instead, dynamic pursuit occupied a population geometry largely shared with ballistic quick captures.

Within this shared geometry, we next asked whether the temporal relationship between preparatory and movement-related activity changed across movement types. Prior work in macaques performing a target jump task suggested that online corrections can reactivate preparatory dimensions while movement-related activity is ongoing, allowing a corrective movement to be planned while the arm is in flight (Ames et al. 2019). We therefore projected pursuit activity into the preparatory and movement subspaces defined during quick capture behavior. Across both marmosets, preparatory-subspace occupancy consistently preceded movement-subspace occupancy for all movement types except corrections (Fig. 3D,i-iv). To quantify this relationship, we defined a preparatory lead index (PLI) as the time by which peak preparatory-subspace occupancy preceded peak movement-subspace occupancy. In both marmosets, preparatory activity led movement activity during initial reaches, whereas PLI was much smaller during corrective movements, consistent with a reduced separation between preparation and execution during online correction (Fig. 3D,v). Reset leads fell between these extremes and differed across subjects, being comparable to initial reaches in TY but substantially smaller in JL. Thus, although pursuit reused the quick-capture geometry, corrective movements altered the timing, and extent, with which preparatory and movement-related dynamics were expressed within that space.

#### Dynamic pursuit increases neural trajectory tangling, especially during corrective movements

Given the obvious need to incorporate ongoing visual feedback of moth position during pursuit of erratically evading prey, we next asked whether dynamic pursuit increased neural trajectory tangling (Q) (Russo et al. 2018). Trajectory tangling asks a simple question: when the neural population reaches a similar state at different moments, does it move in the same direction afterward? Low tangling indicates a consistent flow through state space, whereas higher tangling indicates that similar states can be redirected along different paths, as expected during feedback-driven correction (Russo et al. 2018). During quick captures, tangling remained low during the main population trajectory associated with reaching, with modest increases during later hand-to-mouth movements (Fig. 3E,i, left). Pursuit episodes showed a different pattern: tangling increased during the local trajectory deviations associated with corrective movements (Fig. 3E,i, right). Across episodes, tangling was significantly elevated during dynamic pursuit relative to quick captures in both marmosets (Fig. 3E,ii). Median tangling was 1.34-fold higher during pursuit than quick capture in TY (KS test, p < 10⁻¹⁰, D = 0.26, Cohen’s d = 0.62, the standardized mean difference) and 1.35-fold higher in JL (p < 10⁻¹⁰, D = 0.27, d = 0.57). We then asked whether this increase was concentrated in specific movement types. Tangling measured at event detection was higher during corrections than during quick-capture initial reaches in both animals (Fig. 3E,iii; Wilcoxon rank-sum: TY median Q = 69.3 vs 45.6, p = 1.6 × 10⁻⁵, rank-sum effect size r = Z/√N = 0.46, n = 67 vs 20; JL median Q = 36.5 vs 27.1, p = 2.0 × 10⁻², r = 0.29, n = 52 vs 12).

To test whether elevated tangling during corrections could be explained by kinematic differences between movement types, we fit subject-specific linear models predicting per-event log₁₀(Q) from movement type while controlling for peak hand velocity, total path length, and reach extent. In TY, the correction-versus-quick-capture coefficient remained strongly positive after kinematic control (β = +0.19, p = 1.9 × 10⁻⁴, n = 157 events) and was robust across baseline and single-covariate model variants. In JL, the same effect was positive at baseline but did not reach independent significance after simultaneous kinematic control (β = +0.11, p = 0.07), consistent with the smaller event count and the weaker within-pursuit movement-type gradient in this animal. Corrections were smaller than quick-capture reaches in both subjects, with median reach extents of 3.0 cm versus 7.4 cm in TY and 2.7 cm versus 7.6 cm in JL. Thus, elevated tangling during corrections cannot be attributed to corrections being larger or faster.

Tangling also followed the expected ordering across movement types. Initial reaches during dynamic pursuit and reach resets showed intermediate tangling between quick captures and corrections (TY median Q: quick-capture initial = 45.6, pursuit initial = 49.1, reset = 55.8, correction = 69.3; JL: 27.1, 35.1, 34.4, 36.5). In TY, resets had significantly higher tangling than quick-capture initial reaches (p = 6.5 × 10⁻³, r = 0.41, Bonferroni-corrected), whereas pursuit initial reaches did not differ significantly from quick-capture initial reaches (p = 0.27, r = 0.16). In JL, the same ordering was present descriptively, although neither secondary contrast reached significance (p = 0.06 for both pursuit initial and reset comparisons), consistent with reduced power in this dataset.

These results show that dynamic pursuit increases neural trajectory tangling relative to quick capture, and that this increase is most pronounced during corrective movements. Thus, online correction does not simply extend the quick-capture trajectory; it transiently alters how population activity flows through the shared state space.

#### Rotational dynamics differ systematically across movement types (jPCA)

Given the elevated neural-trajectory tangling observed during corrective movements, we asked whether the movement types composing dynamic pursuit differed in their rotational dynamics. We applied jPCA to each movement type, projecting per-event population activity into the best-fit jPCA plane for that movement class (Churchland et al. 2012). This analysis revealed rotational trajectories across movement types but their rate and coherence varied systematically (Fig. 3F,i-iv). Quick-capture reaches showed the fastest and most coherent rotations. By contrast, online corrections rotated more slowly and showed weaker rotational dominance. Vacillations were the slowest and least rotational. The pattern was consistent across both subjects:

| Type | TY (n, f Hz, R<sup>2</sup>-ratio, plane var) | JL (n, f Hz, R<sup>2</sup>-ratio, plane var) |
| --- | --- | --- |
| Quick-Capture (QC) | 32, 1.17, 0.93, 28% | 14, 0.88, 0.88, 28% |
| Dynamic-Pursuit initiation (DP) | 31, 0.74, 0.79, 29% | 21, 0.84, 0.81, 25% |
| Reset | 39, 0.91, 0.84, 24% | 22, 0.97, 0.86, 22% |
| Vacillation | 20, 0.41, 0.72, 21% | 9, 0.23, 0.61, 20% |
| Correction | 88, 0.79, 0.81, 22% | 58, 0.50, 0.77, 17% |

*f* is the imaginary part of the dominant skew-symmetric eigenvalue expressed in Hz; the R²-ratio is R²(M_skew)/R²(M_best), a rotational-dominance index; plane var is the variance captured by jPC1–jPC2. Each row is a separate jPCA fit on that movement type alone, in the type’s own best-fit plane, the per-type fits drawn as the Fig. 3F,i-iv portraits, not the pooled cross-type fit. JL Vacillation rests on n = 9 events, a small sample reflecting the natural rarity of this movement type; its severely slowed rotation nonetheless matches the slowing seen in the larger TY sample.

A representative pursuit episode from JL, projected into the episode-scale jPCA basis, illustrates this within-episode structure (Fig. 3F,v, left). Large, smooth rotations corresponded to initial reaches and reach resets, whereas smaller, lower-amplitude deviations corresponded to mid-reach corrections. The jPC1 and jPC2 time series show the same structure unfolded over time, with movement-event detections marked beneath the trajectory (Fig. 3F,v, right).

Corrections express slower and smaller rotational dynamics than ballistic reaches. We next asked whether the rotational dynamics observed across movement types differed systematically in frequency, amplitude, or coherence. Across the per-type jPCA fits, the dominant rotational frequency separated quick captures from corrections in both marmosets, with quick-capture rotations faster than correction rotations (Fig. 3F,vi). Because events within the same pursuit episode are not independent, we tested phase-rate differences with a linear mixed-effects model (movement type fixed; subject and episode random), which showed a strong effect of movement type (omnibus p = 5 × 10⁻¹⁹) and significantly slower corrections than quick captures (−0.506 Hz, p = 2 × 10⁻⁷), consistent with an episode-block bootstrap in which the difference excluded zero (95% CI, +0.35 to +0.69 Hz). Per-subject descriptive values and conventions are given on the panels (Fig. 3F,vi) and in the Statistics section of Methods.

Rotational frequency and rotational coherence are distinct axes, and we report them separately. Corrections were reliably slower than quick captures in both subjects, whereas reduced rotational dominance ratio was evident in TY but not in JL. In JL correction dynamics rotated at roughly half the quick capture frequency (0.50 vs 0.88 Hz) while maintaining a similar rotational-dominance ratio. Thus, the main cross-subject difference was not the loss of rotational structure but a shift toward slower correction-related rotations. The reduced rotational-dominance ratio in TY indicates an additional subject-specific reduction in rotational coherence.

We also tested whether the slower phase rate during corrections could be explained by projection into the shared jPCA plane rather than by a genuine difference in correction dynamics. This was not the case. When corrections were projected into their own best-fit jPCA planes, they remained slower than quick captures, pursuit initial reaches, and resets in the same subject. Median own-plane correction frequencies were 0.82 Hz in TY and 0.50 Hz in JL. In TY, the correction plane was closely aligned with the shared plane, with principal angles of 14° and 18°. In JL, the correction plane was more moderately tilted, with principal angles of 32° and 48°, but the shared-plane estimate only modestly under-reported correction frequency. This projection effect was small relative to the approximately 0.4–0.5 Hz difference between quick captures and corrections. Thus, the slower correction dynamics cannot be explained by the choice of projection plane.

Corrections also produced smaller rotational excursions than quick-capture reaches in both subjects (Fig. 3F,vii), confirmed by the mixed-effects model (−0.452 a.u., p = 1 × 10⁻⁹; episode-block bootstrap sign-concordant). Corrections therefore differed from ballistic quick captures along two independent per-event metrics: they were both slower and smaller in rotational amplitude.

Together with the subspace-occupancy and tangling analyses, these results support a distinction between ballistic and corrective dynamical regimes during prey capture. Quick captures, pursuit initial reaches, and resets were associated with larger and faster rotations, consistent with movements initiated from a more canonical premovement state. Corrections, in contrast, occurred while the arm was already extended and while visual feedback continued to update the motor goal. These events showed minimal preparatory lead, elevated trajectory tangling, and slower, smaller rotations initiated from non-canonical state-space positions. Thus, corrective movements did not abolish rotational dynamics; they re-expressed them in a distinct, feedback-driven regime suited to online control.

### A shared sequential firing-rate backbone spans prey-capture movement types

We next evaluated how population-level dynamics corresponded to the firing rate profiles of individual units aligned to each movement type. For every kinematically-detected movement-onset event (Methods, Per-event segmentation), we computed each unit’s peri-event time histogram (PETH) in a detection-centered window from −1.5, +1.5s and normalized each unit to its own maximum firing rate across the five movement types (Methods, PETH computation and normalization). Units were then ordered by their latency to peak firing during full reach onsets, using a single latency value computed from a PETH that pooled all full-reach events across prey-capture episodes (Methods, Latency-only ordering). The same unit order was used for all movement-type columns (Fig. 4B). This single latency ordering revealed a clear sequential activation pattern during prey capture. In quick captures and dynamic-pursuit initial reaches, activity progressed across the population from the period preceding hand extension through extension and into the post-extension hand-to-mouth interval (Fig. 4B). To characterize this structure during quick captures, we classified each unit according to the window in which its mean normalized firing rate was largest: Pre, −1.5 to −0.4 s; Extension, −0.4 to 0 s; or Post, 0 to +1.5 s (Methods, Three-window response classification). In TY, this yielded three temporal response classes — early-tonic (Pre-dominant), phasic-extension (Extension-dominant), and late-tonic (Post-dominant) — in closely matched proportions across subjects (per-class counts and percentages in the Fig. 4 caption). Early-tonic units showed elevated activity prior to hand extension, phasic-extension units activated transiently around movement onset, and late-tonic units showed sustained activity during hand retraction and hand-to-mouth movements.

**Fig. 4.**
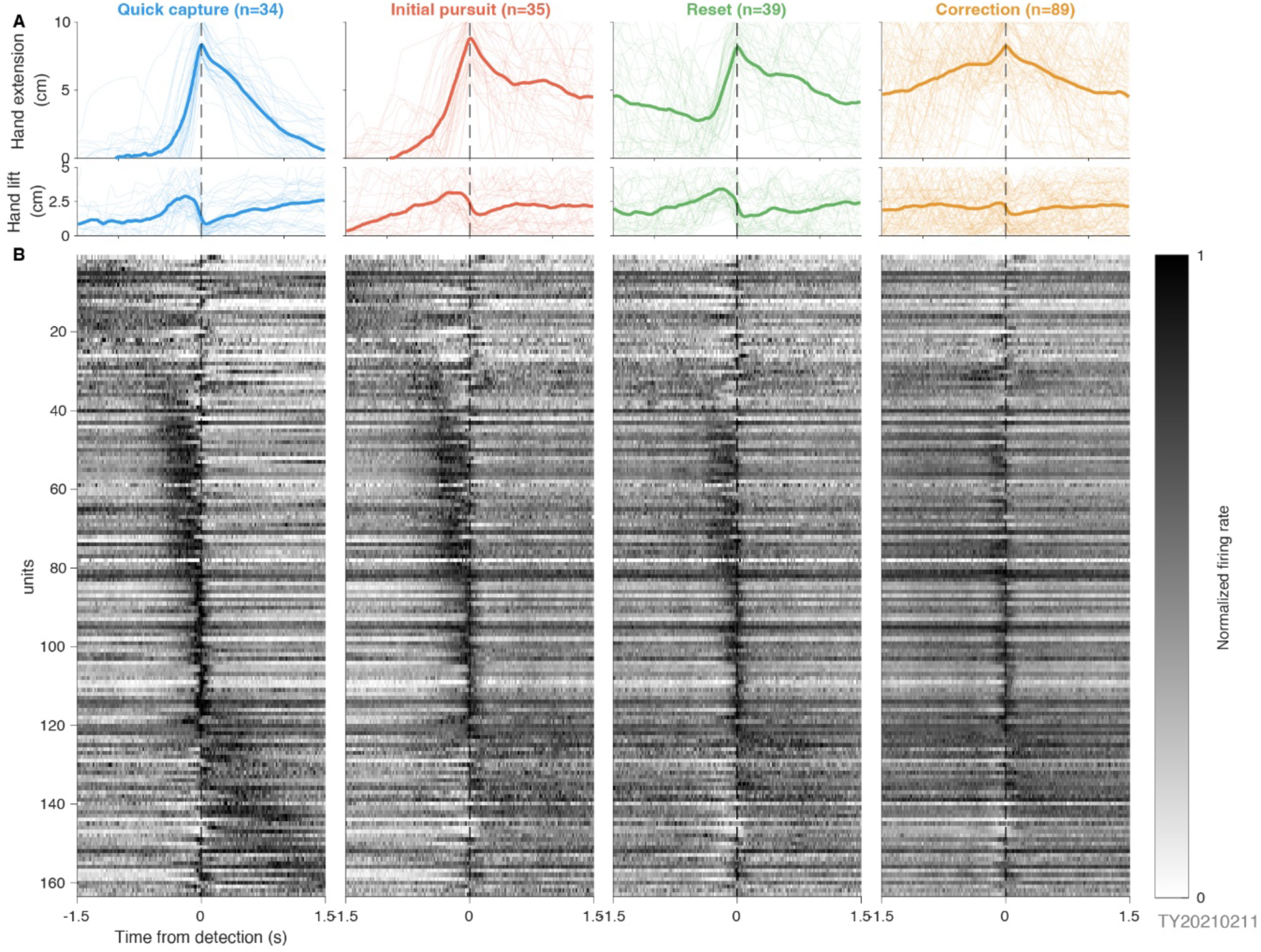
Common backbone of sequential population activity underlies reaching during prey capture. **A)** Per-movement-type mean hand kinematics (extension and lift, cm) for each event recorded during a single session of prey capture in marmoset TY. Mean traces are drawn over individual per-event traces, which are in thinner, lighter color. Movement types (column order): quick-capture initiation (QC), pursuit initiation (DP), reset after withdrawing the hand (Reset), online correction (Correction). **B)** Per-unit peri-event time histograms (PETHs) for each movement type, in the same column order as A. Each row is one unit (TY n=163); columns share the row order. Each unit is normalized to its own maximum firing rate across all five movement types (QC, DP, Vacillation, Reset, Correction even though Vacillation is not displayed). Units are ordered by each unit’s reach-onset latency to peak firing (Methods, Latency-only ordering). Time window: detection-centered, [-1.5, +1.5] s. Detections are roughly peak extension events, see Methods for movement type specific details. Bin size 20 ms; no additional smoothing. Units were assigned to one temporal response class by the timing of their peak quick-capture firing. In TY (n = 163), this yielded 27 early-tonic (Pre-dominant, 17%), 90 phasic-extension (Extension-dominant, 55%), and 46 late-tonic (Post-dominant, 28%) units; JL (n = 133) showed closely matched proportions, with 18 early-tonic (14%), 79 phasic-extension (59%), and 36 late-tonic (27%) units.

Correction-related activity was embedded in this same sequential structure but was generally reduced in amplitude. Early- and late-tonic units, and a subset of phasic-extension units, were less active during corrective movements than during quick captures (Fig. 4B, Correction column). No unit was selectively strongest during corrections; every unit that responded during corrections responded more strongly during at least one other movement type. To quantify this effect by response class, we compared each unit’s mean normalized firing rate in its dominant quick-capture window with its activity in the corresponding window for corrections, resets, and dynamic-pursuit initial reaches (Methods, Class-selective firing-rate contrast). Positive values indicate stronger firing during quick captures than during the comparison movement type; per-class contrast values, CIs, and tests are reported in Results and the pooled class-selective contrast table in the Statistics section (per-class breakdown in Supplementary Table S1).

Corrections produced the largest reduction in firing relative to quick captures, consistent across response classes and subjects: firing was higher during quick captures than corrections in both TY (+15%, 95% CI [+11%, +17%], paired Wilcoxon p = 2.9 × 10⁻¹⁶) and JL (+20%, [+16%, +24%], p = 1.6 × 10⁻¹³). The reduction was graded and correction-specific: resets fell between corrections and full initial reaches, while dynamic-pursuit initial reaches were essentially indistinguishable from quick captures, with one exception, the late-tonic class, which retained a quick-capture-specific post-reach plateau in both subjects. Per-class contrast values, CIs, and tests are given in Supplementary Table S1.

Although corrections showed an overall reduction in firing, the latency-sorted Correction column contained a visible band of activated units. We therefore defined a correction-engaged subpopulation using a rule anchored to the correction-detection-centered PETH (Methods, Correction-anchored peri-event partition). A unit was classified as engaged if its Correction-PETH peak occurred within ±200 ms of correction detection and its quick-capture initial-reach peri-event peak exceeded 40% of its normalized maximum. A unit was classified as disengaged if its Correction-PETH peak occurred within this same ±200 ms window but its quick-capture peri-event peak was below threshold. All remaining units were classified as off-band. In TY, 112 of 163 units were engaged (69%), 4 were disengaged (2%), and 47 were off-band (29%). In JL, 56 of 133 units were engaged (42%), 4 were disengaged (3%), and 73 were off-band (55%). Thus, very few units were active near correction detection without also carrying substantial activity during quick-capture reaches. The main subject difference was instead in timing: TY contained more temporally locked correction-engaged units, whereas JL contained more units whose correction-related activity peaked outside the ±200 ms detection window.

This correction-related recruitment was distributed across the quick-capture sequence rather than confined to a distinct response phase. When units were ordered by Correction-PETH peak time, engaged units formed a contiguous mid-sort block, as expected because both the classification rule and the ordering were anchored to correction timing. However, when the same units were ordered by quick-capture initial-reach peak time, engaged units interleaved with off-band units across the sequential firing-rate cascade (engaged-class row span 162 of 163 in TY, 127 of 133 in JL; Statistics, Pursuit-band partition counts). This indicates that correction-related recruitment is distributed across the population’s quick-capture timing structure rather than concentrated in a separate correction-specific subpopulation.

Collectively, these results show that all prey-capture movement types share a common sequential firing-rate backbone, consistent with the high cross-variance between quick-capture and pursuit subspaces (Fig. 3C,i-ii). During corrections, this backbone is re-engaged at lower amplitude and with altered timing, rather than replaced by a distinct firing-rate sequence. This single-unit organization parallels the population-level results: dynamic pursuit reuses the neural structure of ballistic capture while modifying it for online correction.

### Ballistic prey capture recruits a somatotopically directed sensorimotor cortical wave

We next asked whether the sequential firing-rate backbone observed during prey capture was spatially organized across the cortical sheet. To localize the recording arrays anatomically and functionally, we performed intracranial microstimulation (ICMS) in quietly resting, unrestrained marmosets, assigning each electrode to a cortical area and muscular projection field by the minimum current that evoked a palpable muscle response (Fig. 5A,i-ii; see Methods). Most electrodes were responsive in both animals: 90 of 96 electrodes in TY, with a mean threshold of 9.1 ± 6.8 µA, and 84 of 96 electrodes in JL, with a mean threshold of 10.0 ± 9.0 µA. These ICMS maps localized the arrays across the frontoparietal sensorimotor sheet, spanning PMd, M1, area 3a, and area 3b.

**Fig. 5.**
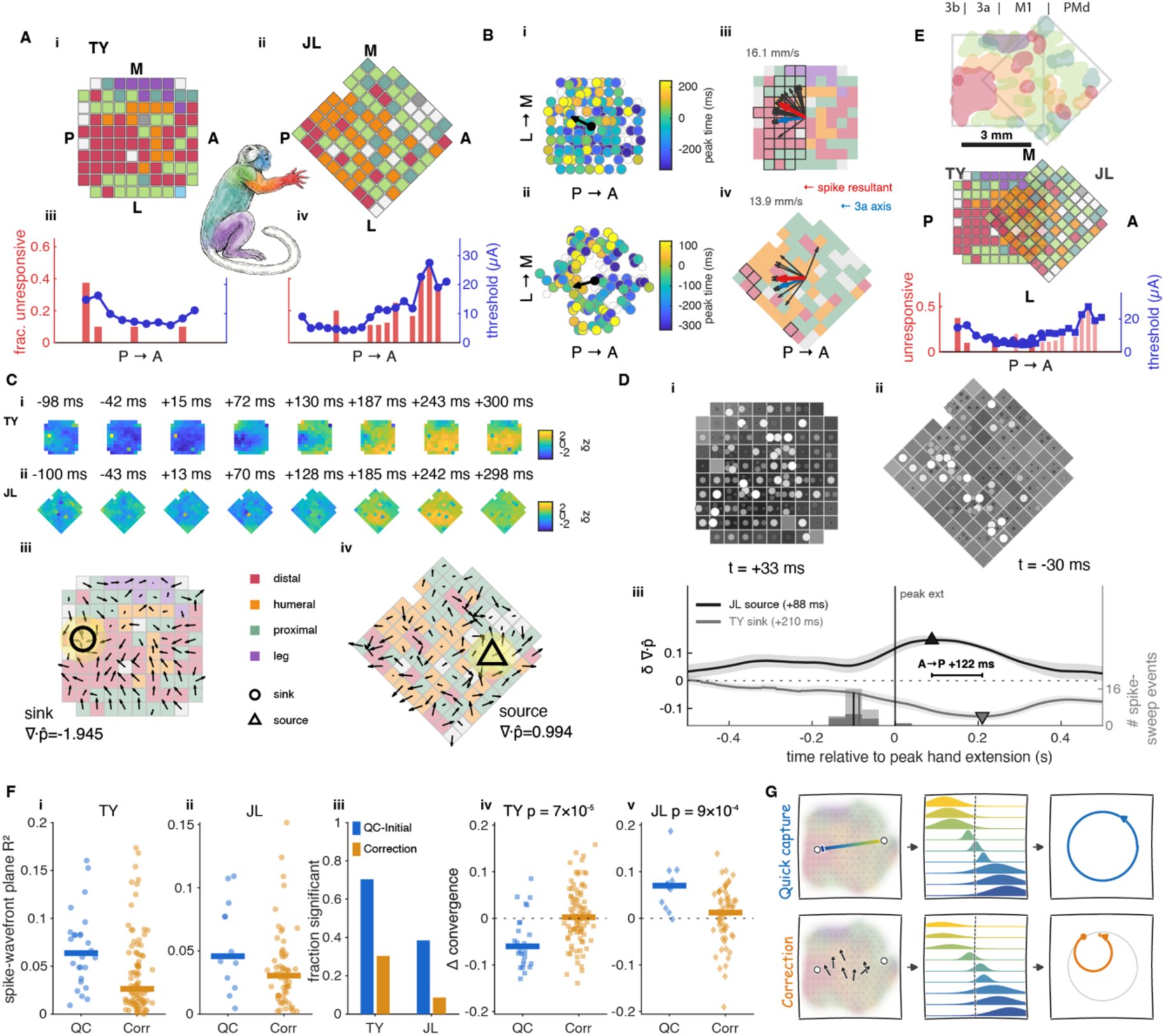
Recruitment-by-somatotopy is a propagating anterior-to-posterior wave aimed at distal forelimb S1, and online corrections disrupt it. **A)** Per-electrode ICMS somatotopy for the two Utah arrays (i, marmoset TY; ii, marmoset JL), colored by the body part in which a muscle twitch was elicited by stimulation. Below each array (iii TY, iv JL), mean ICMS current threshold and fraction of unresponsive electrodes averaged within mediolateral electrode columns along the posterior-to-anterior (P→A) axis; the low-threshold trough locates the most direct descending outflow and anchors the area assignments across the 3a–M1–PMd axis. **B)** Spiking wavefront during quick-capture initial reaches (TY top row, JL bottom row). Per-event planar fits to per-unit firing-rate peak times recover a consistent anterior-to-posterior sweep: (i, ii) single-event spike-recruitment exemplars: one circle per unit at its own array position, filled by that unit’s firing-rate peak time on that single event, with units excluded by the wavefront criterion shown as open circles and the arrow giving the best-fit sweep direction (see Methods, Spike wavefront); (iii, iv) per-event spike-sweep directions for quick-capture initial reaches (thin black arrows; TY n = 27 events, JL n = 13) with their resultant (thick red arrow), shown against the array-centre→area-3a axis defined independently by the ICMS map (blue arrow); the colored tiles behind the arrows are the ICMS somatotopy map of A and C, on the same body-part key; the value printed above each map is the median per-episode planar-fit propagation speed over significant quick-capture episodes (TY 16.1 mm/s, JL 13.9 mm/s). See Fig. S1 for per-event propagation speed and direction distributions. **C)** Co-traveling δ-band (1–4 Hz) field. (i, ii) Single-trial δ filmstrips across peak hand extension (TY, JL) show a field traveling roughly anterior-to-posterior across the array. Phase-flow divergence at the focus shows opposite cross-subject polarities (iii TY convergent sink, ∇·p^ = −1.95; iv JL divergent source, ∇·p^ = +0.99), the two ends expected when one anterior-to-posterior wave is sampled over its destination territory (TY, posterior/3a) versus its launch zone (JL, anterior/PMd). **D)** Spike-to-field correspondence and the implied full-span transit. (i, ii) Single-trial overlays of δ-LFP (greyscale squares, one per electrode, dark = trough) and firing rate (circle brightness; white = high) near peak extension. Every square carries a measured δ-LFP value from its own electrode; no values are interpolated or spatially smoothed (TY +33 ms, JL −30 ms; see Supplementary Video S2, single-trial spike–δ overlays). (iii) On a shared clock anchored to peak hand extension, the δ extrema order JL source (+88 ms) before TY sink (+210 ms), an anterior-to-posterior transit of +122 ms; at the ∼2 mm inter-array centre offset this implies a transit speed near 16 mm/s, which is aligned with the within-array speed band. (This δ-stitch transit is a distinct estimate from the per-episode planar-fit speeds in B.) **E)** Schematic: proposed functional overlap of the two arrays across subjects, registered to the ICMS threshold trough, sampling offset overlapping arms of one premotor-to-3a crescent on the cortical sheet (3 mm scale). **F)** Corrections disrupt the propagation. Per-event spike-wavefront planarity (plane-fit R²), quick captures versus corrections (i TY, ii JL); fraction of individually significant sweeps, quick-capture initial reaches versus corrections, falls TY 0.70→0.30, JL 0.38→0.09 (iii); event-locked change in δ-field convergence around movement detection (iv TY, v JL), one point per event. ∇·p^ is the divergence of the δ phase-flow field — negative where flow converges on a sink, positive where it emerges from a source — and Δ(∇·p^) is its change across movement detection. Quick captures drive each subject’s field further toward its own extremum, so the sign is opposite by construction: TY’s sink deepens (Δ negative) and JL’s source strengthens (Δ positive), while corrections stay near zero in both (rank-sum p = 6.9×10⁻⁵ TY; p = 9×10⁻⁴ JL). **G)** Interpretive sketch (top row, ballistic quick captures; bottom row, corrections). For quick captures, a propagating recruitment wave across spatially ordered cortex produces sequential unit recruitment, which traces a rotation in the dominant population plane. For corrections, recruitment is less coherent and more variably sequenced, tracing a smaller, slower rotational trajectory initiated from a noncanonical region of the population subspace. The sketch depicts the spatial organization underlying the recruitment sequence and its rotational signature.

The two arrays sampled overlapping but distinct portions of this sensorimotor axis. TY’s array extended posteriorly into areas 3a and 3b and contained a prominent distal forelimb representation in its posterior half, flanked by more proximal representations. JL’s array was rotated relative to the cardinal cortical axes and sampled more anterior cortex, with a larger PMd representation and more limited posterior 3a coverage. Across subjects, current thresholds and the fraction of unresponsive electrodes identified a trough of more direct descending output consistent with the expected 3a–M1 axis (Fig. 5A,iii-iv), with TY sampling the posterior 3a–M1 portion of the map and JL sampling the more anterior M1–PMd portion (Fig. 5E). These ICMS-derived maps allowed us to ask whether prey-capture recruitment follows a spatial trajectory across sensorimotor cortex, rather than appearing only as a latency-ordered sequence of individual unit responses.

We therefore tested whether quick-capture reaches were accompanied by a propagating wave of spiking activity across the array. For each quick-capture episode, we fit a planar wavefront to the timing of population recruitment (see Methods); single-event exemplars are shown in Fig. 5B,i-ii. Despite modest single-trial fit values, propagation direction was highly consistent across episodes. Significant anterior-to-posterior sweeps were detected in 19 of 28 quick-capture episodes in TY and 9 of 13 quick-capture episodes in JL, with comparable median per-episode planar-fit propagation speeds of 16.1 mm/s in TY and 13.9 mm/s in JL; the corresponding per-event direction and speed distributions for quick-capture initial reaches are shown in Fig. 5B,iii-iv and Fig. S1. Propagation directions were also consistent across subjects, oriented on the posterior side of the array in both animals. An independent anterior-versus-posterior firing-rate cross-correlation confirmed this directionality, with anterior cortex leading posterior cortex in both TY and JL. Beyond being anatomically posterior-directed, the propagation was aligned with the independently mapped distal forelimb representation. For each event, we projected the propagation direction onto the axis from the array center toward the ICMS-defined area 3a/distal forelimb region (blue axis, Fig. 5B,iii-iv). Propagation directions showed positive projections onto this axis in both subjects, with median projections of 0.83 in TY and 0.95 in JL (the projection is the cosine of the angle between an event’s propagation direction and this axis, so 1 indicates propagation directly toward the distal forelimb representation and 0 indicates propagation perpendicular to it). In TY, all analyzed events projected toward the distal half of the map, and in JL, 12 of 13 events did so. Thus, ballistic quick-capture reaches were accompanied by a coordinated anterior-to-posterior spike wave directed toward the distal forelimb portion of the sensorimotor map. Short-train ICMS tags each electrode by the body part it most readily drives, providing a coarse coordinate for localizing a distal-forelimb territory; we use the map in this sense only, and do not read it as evidence that the underlying cortex is a modular, single-body-part layout.

We next asked whether local field activity provided an independent signature of the same propagating event. In the δ band (1-4 Hz), single-trial field snapshots during hand extension showed a traveling field across the array (Fig. 5C,i-ii). To characterize the geometry of this field, we computed the divergence of the δ phase-flow field, which distinguishes local convergence, corresponding to a sink, from local divergence, corresponding to a source. TY showed a convergent sink during the event-locked reach window, whereas JL showed a divergent source (Fig. 5C,iii-iv). This cross-subject polarity reversal matched the anatomical placement of the arrays: TY sampled the posterior, somatosensory-biased end of the sensorimotor sheet, where the wave arrived, whereas JL sampled the more anterior, premotor-biased end, where the wave emerged (Fig. 5E). Consistent with this interpretation, TY’s δ sink colocalized with distal forelimb cortex in most events, and JL’s δ flow was directed toward the posterior 3a corner of the array. At the single-trial level the two signals (i.e., the local field potentials and spikes) were spatially coincident: in example events near peak hand extension, the units firing most strongly sat over the electrodes where the δ field was at its trough (Fig. 5D,i-ii; Supplementary Video S2). The timing of the spike and field signals further supported a common anterior-to-posterior propagation. On a shared behavioral clock aligned to peak hand extension, the δ extremum occurred shortly after peak extension in both subjects, whereas the spike sweep preceded the δ extremum by approximately 87 ms in TY and 100 ms in JL (Fig. 5D,iii). The ordering of the field extrema across arrays was also consistent with anterior-to-posterior travel: JL’s source preceded TY’s sink by 122 ms (Fig. 5D,iii). Given the approximately 2 mm offset between array centers, this timing implies a cross-array transit speed near 16 mm/s, matching the within-array spike-wave speed and indicating that the same propagation is resolvable across anatomically offset arrays in different animals. Because organized δ propagation did not precede the spike sweep, we interpret the δ signal as a co-traveling field signature rather than as evidence that the field organizes or drives the spike wave.

Finally, we asked whether this propagation differed across movement types. Relative to quick-capture reaches, corrective movements showed a less coherent spike wavefront. Per-event plane-fit R² decreased during corrections in both subjects (Fig. 5F,i-ii), and the fraction of individually significant sweeps fell from 0.70 to 0.30 in TY and from 0.38 to 0.09 in JL (Fig. 5F,iii). Thus, the reduced propagation during corrections was evident at the single-event level and cannot be explained solely by across-trial averaging or phase cancellation from more variable correction kinematics.

The δ-field signature showed the same movement-type specificity. During quick-capture reaches, the event-locked δ field moved toward its end-of-axis extremum: convergence deepened in TY and source strength increased in JL. During corrections, by contrast, the event-locked δ-field change was flat in both subjects (Fig. 5F,iv-v). In TY, quick captures produced a significant increase in convergence, whereas corrections showed no comparable change; the quick-capture versus correction contrast was significant by rank-sum test. JL replicated the same contrast with the opposite source polarity. These differences survived controls for peak movement speed and correction isolation. They also were not explained by where corrections occurred within pursuit episodes, because even early corrections remained flatter than quick-capture reaches.

In sum, these results show that ballistic prey-capture reaches recruit a coordinated anterior-to-posterior wave across sensorimotor cortex, directed toward the distal forelimb representation and accompanied by a co-traveling δ-band field signature. During online corrections, this propagation persists in weakened form: it is less planar, less often significant, and lacks the event-locked field change observed during ballistic reaches. This provides a spatial counterpart to the preceding population analyses: the canonical quick-capture dynamics are expressed as an organized cortical recruitment wave, whereas feedback-driven corrections engage a weaker and less spatio-temporally structured version of the same propagation (Fig. 5G).

## Discussion

Canonical motor-cortical dynamics have been defined largely in trained primates performing constrained reaches, leaving open whether the same principles organize natural actions that are self-initiated, feedback-rich, and continuously corrected. By recording sensorimotor cortex while freely moving marmosets captured evasive prey, we asked whether the canonical dynamical signatures established in constrained reaching generalize to natural primate behavior, and how those dynamics change when movements require online correction.

We found that the major population-dynamical signatures of primate motor cortex persist during natural prey capture. Beyond being present, these signatures were flexibly reconfigured: the same canonical structure accommodated online correction (via slower, smaller neural trajectory rotations, elevated tangling, and compressed or absent preparatory leads) without the population leaving its shared low-dimensional manifold. Quick captures, in which marmosets rapidly reached to seize prey before it escaped, expressed near-orthogonal preparatory and movement-related subspaces (Elsayed et al. 2016) and strong rotational population dynamics (Churchland et al. 2012), despite the absence of an instructed delay. Thus, these signatures are not simply products of delayed reaching paradigms or highly structured laboratory tasks. They can emerge during voluntary, ethologically grounded movements in which the animal initiates action as soon as the prey becomes reachable. This extends the computation-through-dynamics framework (Vyas et al. 2020) from trained reaching tasks to natural primate action.

The added behavioral complexity of pursuit did not produce a wholesale expansion into new neural dimensions. Dynamic pursuit included resets and mid-reach corrections, and the corresponding neural trajectories were more elaborate than those observed during quick captures. Yet these trajectories remained largely within the same low-dimensional geometry. This finding extends conserved-manifold accounts of motor control (Gallego et al. 2017, 2018) to voluntary natural action; adaptations of an existing behavior explore new regions of an already-established subspace rather than building a new one (Perich et al. 2024). Complex pursuit was not expressed primarily by recruiting a separate population state space, but by traversing a shared geometry in different ways as behavioral demands changed.

Within this shared geometry, corrective movements formed a distinct operating regime. During quick captures, pursuit initiations, and resets, preparatory-subspace occupancy preceded movement-subspace occupancy, consistent with a premovement state preceding the subsequent reach. During corrections, this temporal separation was minimal or absent, mirroring the parallel preparation-and-movement geometry reported for mid-reach corrections in constrained primate reaching (Lara et al. 2018; Stavisky et al. 2017; Ames et al. 2019). Corrections also showed elevated neural trajectory tangling, indicating that similar neural states were more likely to be followed by different future trajectories. This is consistent with ongoing feedback or changing task demands redirecting population activity during movement; tangling has been read as an index of input-driveness in dexterous grasping and in feedback-driven manual correction (Suresh et al. 2020; Perich et al. 2024), and as a constraint that shapes which solutions networks express (Saxena et al. 2022), although tangling alone does not identify the source of that input. Because corrections were smaller and slower than quick-capture reaches, the correction-associated increase in tangling cannot be explained by corrections being larger or faster.

The rotational analyses support the same distinction. Corrective movements retained rotational structure, but the rotations were slower and smaller than those associated with ballistic reach initiation. Thus, online correction did not eliminate rotational dynamics. Instead, corrections re-expressed rotational structure from a different region of state space and on a different temporal scale. A comparable organization has been described in motor cortex when a rotational trajectory is executed at different rates: individual-speed trajectories retain their elliptical form but are separated in state space, and artificially rescaling a single trajectory in time without that separation raising trajectory tangling (Saxena et al. 2022).This distinction is important for interpreting the relationship between autonomous dynamics and feedback control. The data do not support either side of a strict dichotomy between motor cortex as a self-contained autonomous generator (Churchland et al. 2012; Russo et al. 2018) and motor cortex as a feedback-driven system; more recent work already qualifies the autonomous picture by showing motor-cortical activity to be substantially input-driven (Sauerbrei et al. 2020) and its rotational structure reproducible by a feedback controller (Kalidindi et al. 2021). Rather, sensorimotor cortex appears to preserve a canonical dynamical scaffold while allowing feedback to redirect, rescale, and retime ongoing activity.

The single-unit firing-rate structure clarifies how this reconfiguration is implemented. Across movement types, units were recruited in a common sequential order, forming a shared firing-rate backbone that spanned quick captures, pursuit initiations, resets, and corrections. Corrections did not recruit a distinct correction-specific subpopulation. Instead, the same population sequence was re-engaged at lower amplitude and with looser temporal locking. This result links the population geometry to the firing-rate data: pursuit modifies a common neural sequence rather than replacing it with a new one.

The spatial analyses provide an additional constraint on this interpretation. The shared sequential backbone was not only an ordering of unit response latencies; it was organized across the cortical sheet. During ballistic quick captures, spiking propagated as an anterior-to-posterior wave directed toward the ICMS-defined distal forelimb representation. A co-traveling δ-band field provided an independent field-level signature of the same propagation. During corrections, this spatial organization weakened: the spike wavefront was less planar, significant sweeps were less frequent, and the event-locked δ-field change was reduced. Thus, the same transition observed in population state space was also visible across cortical space. Ballistic reaches engaged a coherent recruitment wave, whereas feedback-associated corrections engaged a weaker and less temporally structured version of that propagation.

This propagation provides a spatial organization for the recruitment sequence, not a circuit mechanism. The present recordings do not identify the laminar, cellular, or synaptic substrate of the wave, and they do not establish that the δ field drives spiking. Indeed, the spike sweep preceded the δ extremum, so we interpret the field as a co-traveling signature of propagation rather than as evidence for a causal field-to-spike interaction. Nor do we use the wave to argue that rotational dynamics require an autonomous generator: a fixed recruitment sequence can itself produce rotational structure in low-dimensional projections (Lebedev et al. 2019). What our data add is that this sequence is physically organized across sensorimotor cortex and changes systematically with movement type. The rotation, firing-rate cascade, and cortical wave therefore provide complementary views of a common recruitment process.

Propagating waves are a well-established feature of motor cortex, but prior motor-cortical waves are largely agnostic or orthogonal to the somatotopic map: the rostro-caudal β-wave carrying target information (Rubino et al. 2006; Takahashi et al. 2015), its causal role in reaction time (Balasubramanian et al. 2020), and the high-γ amplitude wave carrying reach kinematics (Liang et al. 2023) are all tied to a fixed anatomical axis rather than to somatotopy (Best et al. 2016; Balasubramanian et al. 2020). Additionally, in marmoset area MT spontaneous traveling waves have been shown to gate perceptual sensitivity (Davis et al. 2020). Our result differs in the frequency band (1–4 Hz, on the reach timescale), anchoring (tied to an independent ICMS somatotopic destination in distal forelimb areas 3a and 3b), and use (a discriminator between ballistic capture and feedback-driven correction), and so supplies the somatotopically anchored, spiking evidence the proximal-to-distal hypothesis required but the β-band record had not delivered (Hatsopoulos et al. 2010). What organizes the sweep remains open: intracortical horizontal fibers conduct at ∼0.1–0.8 m/s, roughly an order of magnitude faster than our ∼15 mm/s transit (Muller et al. 2018), while a sequenced thalamocortical drive (Logiaco et al. 2021) or a recurrent network with no dedicated conduction substrate (Luczak et al. 2007) would each fit the slow, reach-matched, feedback-degraded speed we observe; our data do not distinguish them. That this wavefront is resolvable at all may owe partly to the marmoset’s lissencephalic cortex, whose unfolded sensorimotor sheet places the somatotopic map within reach of a single planar array.

These findings support a view of sensorimotor cortex as a feedback-coupled dynamical controller. When the animal is poised and time permits, as in quick captures or reach resets, population activity occupies a preparatory state before movement and then unfolds through a large, fast rotational trajectory. When the prey evades and the arm is already extended, corrections are generated without returning to the canonical preparatory state. In that regime, activity becomes more tangled, rotations become slower and smaller, and cortical propagation becomes less coherent. The moth’s evasive maneuvers therefore reveal a natural analogue of the perturbations and target jumps used to study feedback integration in constrained primate tasks (Stavisky et al. 2017; Ames et al. 2019), but in a continuous behavior where preparation, execution, and correction are interleaved.

By bringing population-geometry analyses to voluntary prey capture, this study shows that canonical motor-cortical dynamics generalize to unconstrained, naturalistic prey capture, but not as rigid templates. Natural action reveals both their stability and their flexibility: a shared low-dimensional structure organizes ballistic capture, while online correction shifts the population into a distinct feedback-associated regime. These results do not settle whether motor cortex is best described as autonomous or feedback-driven, but they tilt the description toward a controller that exploits autonomous dynamics rather than a purely autonomous one: the population state reflects an initial condition and an autonomous trajectory when time permits, and re-expresses those same dynamics under feedback, rather than replanning from rest, when an evasive target forces correction. Framed this way, the canonical dynamical signatures of motor cortex are best understood not as evidence for autonomy per se, but as the reusable scaffold a feedback controller reshapes to meet the demands of natural, closed-loop action.

## Methods

### Data Collection

#### Behavioral recording setup

##### Voluntary behavioral training

This study focused on voluntary natural behaviors; no restraint and no food or water restriction were used to motivate behavior. Marmosets engaged in prey capture voluntarily using a custom-built, modular apparatus mounted to the top of the home enclosure, following the semiautomated voluntary-training approach designed and established by (Walker et al. 2020). Because neither the apparatus nor the wireless neural recording system imposed tethering or restraint, animals entered the apparatus on their own initiative and engaged with it throughout their waking hours, interleaving prey capture with ongoing spontaneous behavior. A prey-capture episode was initiated by dispensing a single moth into the apparatus while the subject was alert, with subsequent prey dispensed after the previous item was captured or escaped.

##### Camera acquisition

Reaching behavior was recorded with high-speed cameras (Blackfly S, 1440 × 1080 resolution; Teledyne FLIR). Two cameras positioned to optimize visibility of the left upper limb recorded marmoset TY at 150 fps, and a multi-camera array recorded marmoset JL at 200 fps. Image acquisition was triggered by the marmoset interrupting an infrared beam-break sensor as it approached the apparatus. Camera exposure times were recorded as TTL pulses on an analog input channel of the Cerebus acquisition system, synchronizing each video frame to the neural recording.

##### Hand kinematics quantification

Three-dimensional hand position was reconstructed from multi-camera video using markerless pose estimation. Two-dimensional keypoints were tracked in each camera view with a DeepLabCut network (ResNet-50; (Mathis et al. 2018), and Anipose was used to perform 2D filtering, 3D calibration, and triangulation (Karashchuk et al. 2021). The triangulated hand traces were post-processed to remove brief tracking lapses prior to analysis.

For the present analyses, kinematics was summarized by a single marker on the left (reaching) hand in both subjects, reported in a workspace coordinate frame. The origin is placed at the marmoset-egocentric left, proximal corner of the prey-capture workspace. The y-axis is aligned with the primary axis of hand extension (away from the marmoset), with the hand beginning near y ≈ 0 at the start of each episode; the z-axis is aligned with gravity (hand lift), with the hand beginning near z ≈ 0; and the x-axis runs along the marmoset-egocentric left–right dimension, with the marmoset positioned at approximately x ≈ 10 cm and the hand at approximately x ≈ 5 cm. Hand velocity [vx, vy, vz] was obtained by differentiating the [x, y, z] hand position. These six variables are the kinematic quantities used throughout.

The behavioral apparatus was designed to elicit primarily reaches with the arm contralateral to the hemisphere in which the electrode array was implanted (left arm for both marmosets). In cases of dynamic prey some bimanual and right hand reaches were performed; however the kinematics of this bimanual behavior was not quantified.

##### Reset correction movement type segmentation

Individual reach events within each prey-capture episode were segmented into movement-type primitives using a custom kinematic algorithm. Candidate movement events were detected from the tracked (left) hand’s 3D position by constructing a composite extension signal that emphasized the reach (Y) axis (composite = 2.0·hand_y + 0.5·|hand_x − x0| + 0.5·hand_z), then identifying local maxima (MATLAB ‘findpeaks’, minimum peak prominence 1.18 cm, minimum inter-peak interval 0.05 s). Peaks occurring before the first reach onset or after the final reach offset, and peaks below a minimum extension of 2.0 cm, were excluded.

Each detected peak was classified by its hand-extension trajectory into one of four movement types: 1) Initial, the first peak reaching full extension (Y ≥ 4.5 cm); 2) Vacillation, an incomplete reach (peak Y < 4.5 cm); 3) Reset, a subsequent peak preceded by a return of the hand toward the body (inter-peak minimum Y < 2.0 cm) following an extended prior reach (prior peak Y > 3.0 cm); 4) Correction, a subsequent peak in which the hand remained extended (no return to the body) between reaches.

The programmatic detections were then manually curated against synchronized video to validate and correct the automatic segmentation: false-positive detections were removed, mislabeled movement types were corrected, and movements missed by the detector (including reaches identified from concurrent population activity) were added. After curation, the event set comprised 217 movement events for TY across all 72 episodes and 124 movement events for JL across all 35 episodes. For event-locked analyses, neural and kinematic data were extracted in fixed windows anchored to each detection (t = 0 at the detected movement).

#### Neural Data Acquisition

##### Surgical Approach

A 96-channel planar Utah array (10 × 10 electrode configuration, 1 mm electrode length, 400 µm inter-electrode spacing; Blackrock Neurotech) was chronically implanted in each marmoset, targeting the forelimb area of primary motor cortex (9.8 mm anterior, 4.5 mm lateral; right hemisphere; JL array rotated 135° relative to TY array), following the surgical approach described in detail by (Walker et al. 2021). The array connector was secured to the skull with a custom, form-fitting titanium pedestal designed to maximize long-term implant biocompatibility and array longevity, replacing conventional bone-cement mounting. Because marmoset sensorimotor cortex is lissencephalic, these areas lie on a smooth, unfolded surface at a spatial scale that matches the array aperture and 400 µm electrode pitch, so a single planar array can sample a functionally meaningful stretch of the somatotopic map, including essentially all of area 3a. In gyrencephalic cortex, such as that of the macaque or human, the equivalent map folds into the central sulcus and would be largely inaccessible to a planar array.

##### Wireless headstage

Neural signals were acquired with a 96-channel CerePlex Exilis wireless headstage (Blackrock Neurotech), used here for both TY and JL. The Exilis houses a digital amplifier and a rechargeable Li-ion battery that supports ∼90 min of continuous recording in a compact package; recording-session length was therefore bounded by battery capacity. The headstage was carried in a custom low-friction housing designed so that the battery/transmitter module could be swapped and recharged without handling or restraining the marmoset via a self-aligning snap-fit connection and a protective helmet over the connector.

##### Neural Recording

Broadband neural signals were transmitted wirelessly from the headstage to a bank of eight receiving antennas, relayed through a digital hub, and acquired by a 128-channel Cerebus system (Blackrock Neurotech). Data were recorded as raw 30 kHz continuous signals and processed offline.

##### Spike sorting

Spike sorting was performed offline on the raw 30 kHz signal using SpikeInterface (Buccino et al. 2020), an open-source framework that wraps multiple spike-sorting algorithms behind a unified interface spanning preprocessing, sorting, cross-sorter comparison, quality-metric computation, and curation. Multiple sorters were run and cross-referenced to retain units identified consistently across algorithms, followed by manual curation. Units were retained at a signal-to-noise ratio threshold of SNR > 3.

Rate estimation. Single-unit firing rates were estimated by binning each unit’s spike train at 1/600 s (≈1.67 ms; 600 Hz) and convolving with a Gaussian kernel of 100 ms standard deviation. Smoothing was applied once, during construction of the dataset, and all downstream analyses used these pre-smoothed rates without further smoothing. For analyses requiring normalization, rates were soft-normalized (dividing by the unit’s firing-rate range plus a constant of 5) following (Churchland et al. 2012).

### Analysis

#### Subspace analyses

##### Subspace identification

Preparatory and movement subspaces were identified by the maxvar joint-orthogonality procedure of (Elsayed’s (Elsayed et al. 2016) maxvar_subspaces), which fits two k=10 dimensional orthogonal subspaces, W_prep and W_move, jointly on the Stiefel manifold to maximize the variance each captures of its respective epoch-specific covariance matrix subject to W_prep ⊥ W_move. Per-event firing rate windows (population rates smoothed with 100 ms gaussian kernel) were drawn from quick capture episodes and soft-normalized (range normalized with a 5 spikes/sec offset, following Elsayed et al. (Elsayed et al. 2016) across all units before fit. The fitting epochs were a preparatory period [-500, -200] ms relative to detected movement onset and an execution period of [-100 ms, peak hand extension]. Rates were mean-centered (no whitening) before projection. The same W_prep / W_move bases were used for the analysis in Fig. 2C,D and Fig. 3D, so all projections derive from a single basis for each subject.

##### Alignment index

The alignment index A between a source subspace and a target condition is the fraction of target-condition variance captured by the source subspace, normalized by the top-k spectral mass of the target covariance: A = trace(Wᵀ C W) / Σᵢ₌₁ᵏ λᵢ(C), where W is the source-subspace basis and C is the target-condition covariance with ordered eigenvalues λᵢ. For Fig. 2C the alignment index and its random-subspace null (Elsayed and Cunningham 2017) were evaluated at k = 10, the full dimensionality of the joint-orthogonal W_prep / W_move bases (captured variance normalized by the top-10 eigenvalues of the target covariance). The scree panels separately report each subspace’s effective dimensionality at an 80%-variance threshold (k_prep = 7 and k_move = 5 in TY; 7 and 4 in JL, a lower-dimensional summary of the same 10-dimensional fit, so the 10-dimensional subspaces capture somewhat more than 80% of their epoch variance) and the cross-projected variance fractions are summed over these thresholded dimensions. For the quick capture v dynamic pursuit comparison in Fig. 3C (top row), per-condition bases were obtained with standard principal component analysis, joint orthogonality is unnecessary here because the two conditions’ subspaces are so well aligned that an orthogonality constraint would obscure their overlap, with k set by a per-condition 90% variance threshold (k = 51 in TY, k = 45 in JL). Observed alignment was tested against a manifold-constrained random-subspace null (following Elsayed, 1,000 draws of k-dimensional random subspaces sampled from a manifold biased to the global quick capture plus dynamic pursuit covariance). P-values are one-tailed empirical percentiles, with the tail chosen by the direction of the claim: a left-tail test for orthogonality (low observed A relative to the null) and a right-tail test for overlap (high observed A relative to null).

##### Preparatory lead index (PLI)

For each (subject, movement-type) combination, occupancy timecourses in W_prep and W_move were computed as the per-event sum of squared projection magnitudes, normalized separately per subspace (the display convention of (Lara et al. 2018), which prevents a scale mismatch between preparatory and movement dimensions from masking the lead structure) and averaged across events. Peak times of the preparatory and movement occupancy curves were located by searching backward from the event-detection moment (t = 0) within the display window [−0.75, +0.50] s, so that when multiple local maxima are present the peak closest to detection is reported. Latency was measured by peak position rather than by a threshold-crossing of each occupancy curve (e.g. the 10%-of-peak onset latency of (Lara et al. 2018)) because the continuous, multi-event structure of naturalistic pursuit affords no quiescent pre-event baseline against which a fixed-fraction threshold could be defined; peak position is well-defined without a baseline, at the cost of greater sensitivity to curve shape for broad or subtly bimodal occupancy traces. A direct cross-check confirmed this choice: a 10%-of-peak onset-latency criterion pinned the onset to the analysis-window edge for most movement types (the occupancy curves seldom fall below 10% of their peak within the window) and was well-defined only for resets, where it agreed with the backward-peak lead (TY +256 ms vs +157 ms; JL +62 ms vs +77 ms).

The PLI is defined as the movement-occupancy peak time minus the preparatory-occupancy peak time, so positive PLI indicates the preparatory subspace peaks first. Peaks with prominence below 0.09 (in display-window-normalized units) are flagged with * in Fig. 3D,iv as modest; the only such flag is JL correction (PLI +72 ms, preparatory-peak prominence ≈ 0.021). JL reset’s preparatory peak (prominence ≈ 0.114) is just above the threshold but visibly smaller than JL’s initial-reach preparatory peaks (≈ 0.41 for dynamic pursuit, 1.00 for quick capture).

Because the PLI measures latencies of tens of milliseconds while the occupancy curves derive from rates pre-smoothed with a 100 ms Gaussian kernel, we verified that the prep-lead pattern is not an artifact of that kernel by re-estimating the continuous rates across Gaussian kernel widths from 50 to 200 ms (bases held fixed) and recomputing the PLI. The qualitative pattern was preserved in both subjects at every kernel width tested: a substantial positive preparatory lead at movement initiations (quick-capture and dynamic-pursuit initial reaches) and a near-zero or negative lead at within-reach corrections, with initiations ordered well above corrections in both subjects. Reset leads were subject-dependent, comparable to initial reaches in TY but substantially smaller in JL, and were the most kernel-sensitive values in the sweep. The initial-minus-correction preparatory-lead gap remained large and sign-consistent at every width (77–173 ms in TY, 143–218 ms in JL). Absolute PLI magnitudes are kernel-dependent, most notably the TY initial reaches, which vary by up to ≈ 100 ms and non-monotonically across kernels, reflecting the sensitivity of any peak-based estimator to fine occupancy structure that a 100 ms kernel smooths over. Narrowing the kernel did not systematically reduce the preparatory leads (they fall for some TY movement types but rise for JL), so the leads are not a by-product of over-smoothing and the near-zero correction lead is preserved or strengthened at narrower kernels. We therefore retain the 100 ms kernel (the most stable peak estimate), treat the per-cell PLI values as approximate latency estimates, and rest the claims on the across-type ordering and the differential prominence of the preparatory peaks rather than on precise millisecond magnitudes.

##### Episode classification

Quick-capture and dynamic-pursuit labels follow the per-episode behavioral coding established for these sessions (TY: 30 quick-capture, 39 dynamic-pursuit, with three non-moth reaches unclassified; JL: 13 quick-capture, 21 dynamic-pursuit after excluding one stationary-prey episode). For both subjects the W_prep / W_move bases were fit on hand-annotated quick-capture episodes. For TY, two right-hand captures were excluded because recording was from the left hemisphere, and one further quick-capture episode was excluded for missing data, leaving 27. This pool was further restricted to isolate pure single-reach captures whose preparatory and execution epochs are unambiguously separable (n = 22): three episodes containing a vacillation event in or adjacent to the reach, one bimanual capture, and one multi-grasp-to-capture sequence were excluded. For JL the full annotated quick-capture pool (n = 13) was used. The preparatory lead index was computed for each of the four movement types, with per-type event counts reported in the Fig. 3D panels.

#### jPCA

##### jPCA fitting

For each movement type, per-event firing-rate windows (250 ms before to 150 ms after movement detection; population rates smoothed once with a 100 ms Gaussian kernel during composite-dataset construction) were reduced to six principal components and fit with jPCA (Churchland et al. 2012), yielding the skew-symmetric dynamics matrix M_skew and its best unconstrained counterpart M_best. Rotational frequency is the imaginary part of the dominant M_skew eigenvalue; rotational dominance is the ratio R²(M_skew)/R²(M_best), which is insensitive to the absolute strength of linear structure and therefore separates how rotational a trajectory is from how predictable it is. Reporting the dominance ratio rather than raw rotational fit quality matters because raw R²(M_skew) is strongly coupled to eigenvalue magnitude (within-type r ≈ +0.9), so a “less coherent” claim taken from raw R² could merely restate “slower”; with the ratio, the apparent JL Correction coherence reduction disappears (confidence intervals overlap) as frequency-coupling artifact leaving the defensible cross-subject claim that Corrections are reliably slower while a coherence reduction is TY-only.

The exemplar portrait (Fig. 3F,v) uses an episode-scale jPCA basis (reaching-window quick-capture and dynamic-pursuit episodes, ±0.5 s buffer) so that the multi-reach, episode-level rotational mode is visible. This basis is used only to display structure; per-event rates are measured in the event-scale basis. The two bases are deliberately not conflated: projecting 400-ms event windows through the episode-scale basis depresses every type’s apparent rate uniformly through a window/period mismatch (0.05–0.56 Hz), a measurement artifact of fitting a short window with a long-period basis rather than biology, which is why rate comparisons are made only within the event-scale basis. The exemplar phase-portrait axes are rendered as an isotropic 1-jPC-a.u. scale bar, since only relative geometry is interpretable.

##### Cross type plane and per-event metrics

The cross-type plane used for the per-event frequency and amplitude comparisons (Fig. 3F,vi-vii) is a single jPCA fit on all five movement types pooled, so the basis is not biased toward any one type’s rotational plane. A per-type best-fit plane is by construction the maximally rotational view of that type, which would make a cross-type comparison performed in per-type planes circular. The cost is that a type whose intrinsic plane is tilted relative to the shared plane is under-reported there, so cross-type differences read in the shared plane are conservative lower bounds on the intrinsic differences (handled by the own-plane control below). Per-type point frequencies and their 95% confidence intervals (Fig. 3F,vi caption) use the bootstrapped dominant M_skew eigenvalue, resampled within type for percentile intervals. This bootstrap is an estimation device for the per-type confidence intervals, not a hypothesis test: a significance test computed on the bootstrap distribution itself is invalid here because the bootstrap-distribution variance shrinks with the resample count rather than reflecting genuine between-group sampling variability, so its p-values collapse to zero artifactually. Inference is therefore carried by the mixed-effects model on the per-event phase rate (below), with an episode-block bootstrap as a non-parametric cross-check. The per-event phase rate is R²-independent by construction, so the slower-versus-less-coherent distinction is not a frequency–coherence coupling artifact.

Amplitude (Fig. 3F,vii) is the per-event detrended peak-to-peak excursion in the jPC1–jPC2 plane: the per-event mean and linear trend are removed and the bounding-box diagonal of the residual loop is taken, making the metric origin-independent and drift-robust. Cross-type comparison is restricted to quick captures, pursuit initiations, resets, and corrections in the shared plane. Throughout, the tilt between a movement type’s own best-fit plane and the shared plane is summarized by their principal angles (two 2-D planes have two principal angles; 0° denotes coincident planes and 90° fully orthogonal ones, so larger angles mean a more tilted plane whose shared-plane projection is more foreshortened). Vacillation is measured in its own intrinsic plane, which sits at a steep tilt to the shared plane (principal angles 78–82°, where a shared-plane projection would foreshorten it), and reported separately, outside the cross-type test. JL Corrections, only moderately tilted (32°, 48°), are measured in the shared plane: a 48° tilt caps cosine foreshortening of apparent amplitude at ≈ 1 − cos 48° ≈ 33%, whereas the observed JL Correction rotational amplitude is roughly half that of quick-capture (median ≈ 0.50× the quick-capture amplitude, a ≈ 50% reduction; Cohen’s d = 2.15). Geometry can therefore account for at most about two-thirds of the observed drop, so the reduction reflects a genuine decrease in rotational excursion, not a subspace-tilt artifact.

To confirm that the slow and small Correction estimates in the shared plane are not subspace-tilt artifacts, Corrections were re-measured in their own best-fit plane and the principal angles between that plane and the shared plane were quantified. Own-plane Correction frequency remained severely slowed (median TY 0.82 Hz, JL 0.50 Hz); TY Corrections were near-aligned with the shared plane (principal angles 14° and 18°) and JL moderately tilted (32° and 48°), with the shared plane only mildly under-reporting JL (0.50 → 0.41 Hz, small against the ≈ 0.4–0.5 Hz gap). The biological effect dominates any projection-geometry term.

##### Existence of rotational structure

Because smooth high-dimensional signals can spuriously appear rotational, the observed R²(M_skew) was compared against a null distribution built from time-shuffled data (the same control used in the Churchland et al.(Churchland et al. 2012) supplementary materials). Observed R²(M_skew) exceeded the time-shuffled null at p < 0.01 for quick capture, dynamic pursuit, reset, and correction in both subjects (Vacillation not assessed, small n and near-orthogonal geometry).

#### Tangling

Neural trajectory tangling (Q) (Russo et al. 2018) quantifies how predictable the future evolution of a neural population trajectory is from its current state, by comparing the velocity vectors at all pairs of population states. For each pair of timepoints (t₁, t₂),

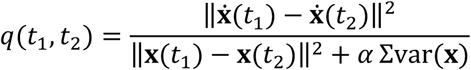

with x(t) the PCA-projected population state, ẋ(t) its time derivative, and α = 0.1 a regularization constant. Q(t) is reported as max over t₂ of q(t, t₂). High Q(t) implies that the state at t is near other states whose subsequent dynamics differ substantially, signaling input-driven (non-autonomous) behavior.

##### Population formatting

For each subject (TY 163 units, JL 133 units) and each condition (Quick Capture or Dynamic Pursuit), we constructed a state matrix by concatenating the pre-smoothed firing rates of all episodes assigned to that condition (manual ground-truth labels). Firing rates were pre-smoothed (100 ms Gaussian kernel, 600 Hz) and not re-smoothed in the tangling pipeline. We applied Churchland-style soft normalization (softenNorm = 5 Hz) and mean-centering, then reduced dimensionality via PCA, retaining numPCs = 15 components per condition. Velocity ẋ was computed as first differences of the PC trajectories divided by samplingRate = 0.01 s. Pairwise q was evaluated over a subsampled grid (timeStep = 5).

##### Within-condition pairing

Q was computed with restricting state pairs to within-episode pairs (treating each episode as a separate “condition”, withinConditionsOnly = true, in the Russo formulation). This follows (Perich et al. 2024), who computed tangling independently per trial for naturalistic single-trial reach-and-pull data and confirmed that this per-trial computation gives results comparable to the global cross-trial computation. Because naturalistic prey-capture episodes lack a common kinematic template that would enable Russo-style cross-condition warping, within-episode computation is the principled choice for our data: cross-episode behavioral variability (posture, reach direction, episode duration) would otherwise inflate Q as a confound rather than as evidence of input-driven dynamics, as our pilot validation confirmed (the global pairing produced a ≈150× inflation in absolute Q scale and an artifactually high DP/QC ratio ≈68× at small timeStep values). With this within-episode formulation, Q at time t indexes whether the population trajectory remains predictable from its own immediate history within the same behavioral episode. It behaves as a measure of within-episode dynamical consistency rather than global state-space autonomy across the whole session. Because the within-condition formulation evaluates Q over a strict subset of the pairs the global formulation would consider, local Q ≤ global Q by construction; demonstrating elevated within-episode Q during pursuit is therefore evidence of input-driven dynamics without requiring the global formulation.

Episode exemplars and trajectory coloring (Fig. 3E,i). Per-condition PCA bases (15 PCs per subject, per condition) were used for the trajectory plots. Episode coloring maps log₁₀(Q) onto a desaturated red-white-blue scale, with the colorscale per subject centered between the QC and DP medians: TY [1.09, 2.09], JL [0.89, 1.89]. Exemplars: JL (V17, E1) (QC) and (V29, E1) (DP).

Distribution-level analysis (Fig. 3E,ii). All Q(t) values produced by Russo’s tangleAnalysis were retained per subject per condition and compared on the log₁₀ scale. n timepoints (TY): 1,440 QC / 4,972 DP. n timepoints (JL): 2,295 QC / 4,850 DP.

Event-level analysis (Fig. 3E,iii). Per-event tangling was sampled from the episode-level Q cache rather than recomputed in event-aligned windows. For each movement type event (TY: 217 valid events; JL: 124 valid events), the event’s detection time on the session clock was converted to a time relative to the onset of the first reach of the episode containing that event, with episodes matched by video and within-video episode index. Q was linearly interpolated from the episode-level cache over a [−250, +150] ms window around detection, and the per-event median was used as the strip-plot value. This approach preserves a single PCA basis and Q scale across Fig. 3E,ii and 3E,iii, enabling direct comparison.

#### PETH heatmaps

##### Per-event segmentation

Movement-onset events were drawn from a manually-curated event set layered on automated kinematic detection (TY: 217 valid events after exclusion of false positives and signal-dropout episodes). Events were partitioned into five movement types, QC-Initial, DP-Initial, Vacillation, Reset, Correction, using each event’s curated movement-type annotation combined with the QC-versus-DP classification of its parent episode. Per-column event counts in Fig. 4 are the post-exclusion totals.

##### PETH computation and normalization

For each movement-type event, we extracted spikes in a detection-centered window of [−1.5, +1.5] s with 20-ms bins and no additional smoothing. Per-unit per-event spike counts were converted to firing rates (Hz) and averaged across events within each movement type. After averaging we applied global per-unit normalization across all five movement types: each unit was divided by its maximum mean firing rate across the concatenation of all five movement-type columns. The Vacillation column participates in the normalization even though it is not displayed in Fig. 4B; this ensures the per-unit maximum of 1 is comparable across units regardless of which column carries each unit’s peak response. Per-unit max-column distribution (TY): QC 52%, Vacillation 24%, Reset 13%, DP 10%, Correction 0%.

##### Latency-only ordering

The unit-row ordering in Fig. 4B is by ascending per-unit latency to peak firing, a scalar derived from a single z-scored, 350 ms (7-bin) moving average PETH aligned to all full-reach onsets pooled across all moth-capture episodes (n=77 pooled onsets in TY). “Full reach onsets” refers to the kinematic event set produced by the first-pass behavior parse where onsets were detected as the preceding hand-speed threshold crossing (5 cm/s) before the hand crossed the apparatus partition into the workspace, with consecutive reaches separated by a minimum of 100 frames and validated against hand-extension peaks (y > 3.85 cm). This set groups initial reaches (in both quick-capture and dynamic-pursuit episodes) with reset reaches (re-extension after retraction) and excludes the mid-reach corrections and vacillations that are annotated in the curated event set (concordant on 77% of TY moth episodes; the remaining 23% show off-by-one or off-by-two boundary differences between the kinematic-parse and curated segmentations). The latency scalar is therefore a per-unit summary of when the population responds across full-reach onsets, used here purely as a one-dimensional ordering of the displayed PETHs; the sort is independent of the movement-type-specific PETH columns it organizes.

The correction-engaged partition (see below) is defined on the column alignment events rather than on this ordering: the band criterion is read on correction detections, the amplitude criterion on quick-capture initial-reach detections. The two alternative row orderings reported in Results (1. by Correction-PETH peak time and 2. by quick-capture initial-reach peak time) are argmax sorts on those column events and are distinct from the Fig. 4B ordering.

##### Three-window response classification

For Results-level characterization, each unit was classified by which of three operationally-defined windows held its largest mean normalized firing rate on the QC PETH column: Pre: [−1.5, −0.4] s → early-tonic; Extension: [−0.4, 0] s → phasic-extension; Post: [0, +1.5] s → late-tonic. The boundaries were set from inspection of the figure heatmap; they are not optimized parameters. Classification is by argmax of the three window means.

##### Class-selective firing-rate contrast

For each unit we computed the mean normalized firing rate in each of the three windows (Pre, Extension, Post) on the QC column and on each comparison column (Correction, Reset, DP-initial), and assigned the unit to a dominant-window class by argmax of its three QC-column means (identical to the three-window response classification above; the classifier is fixed by QC so per-class n is constant across contrasts). For each contrast (QC vs Correction, QC vs Reset, QC vs DP-initial) and each class, we tested whether the QC and reference window-means differed using a paired Wilcoxon signed-rank test (per-class n: TY 27 / 90 / 46; JL 18 / 79 / 36); 95% CIs on the median paired difference (and on its expression as a percent reduction relative to median QC) were obtained by 2000-iteration paired-median bootstrap (unit indices resampled with replacement; seed fixed for reproducibility). The “population pooled” row reports the same contrast with each unit contributing the absolute mean from its own dominant window (per-class union).

##### Correction-anchored peri-event partition

The pursuit-subpopulation partition cited in Results was computed from two columns of the same five-movement-type PETH set used elsewhere in this section (analysis window [−1.5, +1.5] s; 20-ms bins), with one additional preprocessing step. For each unit and each of the QC-Initial and Correction columns we applied a Gaussian smoothing kernel spanning a 7-bin window (≈27 ms standard deviation at the 20 ms bin size) to the normalized PETH; let QC_sm(i, t) and CR_sm(i, t) denote the smoothed traces. We then computed two per-unit scalars:

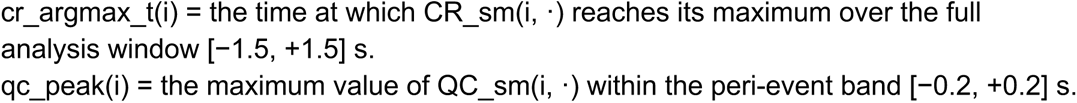

A unit is engaged if cr_argmax_t ∈ [−0.2, +0.2] s AND qc_peak > 0.40; disengaged if cr_argmax_t ∈ [−0.2, +0.2] s AND qc_peak ≤ 0.40; off-band otherwise. The 400-ms band centered on the detection event is event-symmetric and matches the visible temporal extent of activation in both subjects on the Fig. 4B heatmap; the 0.40 amplitude threshold is a lower-tail cutoff on the per-unit qc_peak distribution, which is unimodal and left-skewed (0.40 lies at the 5th percentile in TY, the 22nd in JL). Varying the threshold across 0.30–0.50 leaves the per-class counts within 5 units in both subjects; widening the band to ±300 ms is the more consequential perturbation, moving the engaged count by 7 (TY 112→119; JL 56→63).

##### Three alignment events

The partition rule and the figure sort live on three different anchors that should not be conflated. (i) The Fig. 4B sort key is the argmax of a single PETH locked to all full-reach onsets pooled across episodes (Methods, Latency-only ordering). (ii) The QC-Initial PETH column is detection-centered to QC-Initial event detection. (iii) The Correction PETH column is detection-centered to Correction event detection. Each column’s t-axis is independent. The correction-anchored partition uses anchors (ii) and (iii); the figure sort uses (i).

##### Sort-versus-classification correspondence

The latency sort key (full-reach-onset PETH) and the three-window classifier (QC column window-mean argmax) emphasize different aspects of each unit’s response. To quantify the correspondence, we binned the per-unit latency scalar into the same three windows and cross-tabulated against the QC classifier: 35% of TY units (57/163) and 31% of JL units (41/133) fell into the same window under both definitions. In TY the largest off-diagonal cell was units whose pooled full-reach PETH peak was Post but whose QC-column dominant window was Extension (n=82). This drift is the structural source of the visible heterogeneity within each latency band in Fig. 4B.

Although the latency-only ordering provides a clean single-axis arrangement of these units, sort position and three-window classification do not correspond one-to-one: 65% of TY units (106/163) and 69% of JL units (92/133) are classified into a different temporal window than the one their full-reach-onset peak time falls into, reflecting that the pooled full-reach PETH emphasizes the population’s post-onset peak across all full reaches (which include resets), while the per-movement-type classifier reads each unit’s quick-capture activity profile specifically.

#### Propagating activity

##### Movement-type scope and event counts

The primary contrast is quick-capture initial reaches (the first ballistic reach of a quick-capture episode) versus corrections (TY: 27 quick-capture-initial versus 89 correction events; JL: 13 versus 58). The ballistic group is restricted to quick-capture-initial reaches; expanding it to all initial reaches (quick-capture plus dynamic-pursuit) is a documented robustness secondary that firms TY (Δ p = 4 × 10⁻⁴) but dilutes the JL δ-change (p = 0.22, while JL’s absolute-POST magnitude survives, p = 0.008), so quick-capture-initial is the primary ballistic scope.

##### δ field and phase-flow

LFP was band-limited to δ (1–4 Hz), the reach timescale (one cycle ≈ 250–1000 ms). At each electrode and time we formed the analytic signal z of the δ-band LFP and took its instantaneous phase φ = arg(z). The phase-flow field is the unit-normalized spatial gradient of this phase, p^ = ∇φ / |∇φ| (with ∇φ = Im(∇z / z)): a unit vector at each array site pointing along the local direction of phase advance (Muller et al. 2016). Two scalars summarize the field’s geometry at the focus of the wave. (i) The spatial divergence of the flow, ∇·p^ = ∂p^ₓ/∂x + ∂p^_y/∂y, is negative where flow converges (a sink, the wave arriving) and positive where it diverges (a source, the wave launching)(Davis et al. 2020). (ii) The phase-gradient directionality (PGD) is the resultant length of the phase-gradient directions taken across the array, near 1 for a planar wave whose gradients all point one way and near 0 for spatially incoherent phase; it indexes how wave-like the field is (Rubino et al. 2006). Event-locked PRE and POST windows were taken around movement detection, and the discriminating quantity is the change in divergence at the focus, Δ(∇·p^) = POST − PRE. Because divergence is a coordinate-free (frame-invariant) scalar, the 135° rotation applied to JL’s display cannot convert a convergence into a divergence or otherwise alter the sink/source sign.

##### Channel quality

No per-channel quality rejection is applied at any stage of the propagation pipeline. To establish that this does not affect the reported δ-field results, we flagged channels whose 1–4 Hz amplitude exceeded five times the array median, or whose maximum correlation with any other channel fell below 0.6, in every one of ten interior 200 s windows spanning the recording (boundary windows were discarded as potentially containing recording-onset and -offset artifacts). This flagged 5 of 96 electrodes in TY and 6 of 96 in JL, none of which carried a sorted unit. Recomputing the δ-derived quantities with these channels excluded left every reported result unchanged: the divergence polarity (convergent sink in TY, divergent source in JL), the quick-capture versus correction Δ(∇·p^) contrast (p < 10⁻³ in both subjects before and after), the cross-array transit (122 ms versus 120 ms), and the spike–δ lead. This analysis is a sensitivity check rather than a claim to a canonical bad-channel criterion.

##### Spike wavefront

Per event, the spike wavefront is a planar fit to the per-unit peak times of the pre-smoothed firing rate (each sorted unit’s smoothed-rate maximum within the event window, placed at that unit’s array position; a unit is included if its in-window maximum reaches at least its own mean + 1 SD, and edge-peaking units are excluded). In the single-event exemplars of Fig. 5B,i-ii, each included unit is drawn as a filled circle at its array position, coloured by its own peak time on that event, with the colour scale spanning the 10th–90th percentile of peak times within that event; units failing the inclusion criterion are drawn as faint open circles, and units sharing an electrode are offset so each remains visible. Per-event propagation significance follows the shuffle test of (Liang et al. 2023): electrode locations are shuffled 500 times and the plane refit on each shuffle, and an event counts as a significant sweep when its planar-fit R² exceeds the top 5% of that shuffle null. The unit layout is built unrotated for TY and rotated 135° for JL so that the spike foci and the somatotopy reference share one anatomical frame. The two quantities not to be confused: per-episode planar-fit R² (low, the single-trial wave is weak) versus the directional resultant R (high, the per-episode propagation directions are concentrated).

##### Transit versus reach duration (pooled estimator)

To test whether the spike-wave transit is fixed (conduction-like) or rescales with reach duration (movement-clocked), we estimated transit in a pooled, gate-free form. Per quick-capture-initial reach, we took the per-unit spike peak-time field, averaged it across events within reach-duration tertiles (and, separately, a sliding duration window), and fit a single plane per bin, whose gradient gives one transit per duration bin. Pooling before fitting is deliberate: gating individual events on planar-fit significance would select on R², which itself tracks reach duration, biasing the very comparison being made. Reach duration is detection minus movement onset, where detection is peak hand extension and onset is back-tracked from the full-episode hand-speed trace at 10% of peak speed. The long-minus-short transit contrast was assessed with a 5000-sample bootstrap. This per-event detection-anchored estimator is distinct from the per-episode planar fit reported for Panel B; the result is exploratory and is reported as a Discussion observation only.

##### Somatotopy colocalization

The distal/3a reference is the independent ICMS somatotopy map (electrode microstimulation, not spike timing): distal electrodes are those whose somatotopy labels fall in the hand/forearm/wrist/finger group, and the 3a set is the electrodes whose units are 3a-classified. Two geometric anchors are derived from this map: the array centre (the centre of the 96-electrode grid) and the 3a centroid (the mean array position of the 3a electrodes). The δ-flow sink is tested against this reference by two complementary readouts: a categorical test of whether the sink falls on a distal-labelled electrode, against the interior electrode-share null (binomial), and a continuous test of the sink-to-distal-centroid distance against a uniform interior-site null (5000 draws). For the spike wave, which is planar and has no single convergence point, a directional test projects the per-event propagation vector onto the array-centre-to-3a-centroid axis (signed-rank versus 0); this directional test carries the spike-pillar result. In TY the per-event direction is strongly positive on this axis (median projection 0.83) but points about 49° medial of the array-centre-to-3a-centroid ray (empirical wavefront median 137°; the wave follows the anatomical anterior-posterior axis while the 3a centroid sits mid-row), so the test localizes the distal half as the destination rather than a pinpoint area-3a target. The array’s posterior edge also bounds the observable destination: TY’s convergence sink localizes to the posterior-medial corner, so we cannot exclude a somatotopic destination further posterior than the array samples.

##### Kinematic clock and cross-subject coregistration

Each reach was anchored at its own peak hand extension. Clock comparability across subjects was established by direct comparison (sharp anchor in both; statistically matched reach extent, duration, and path length; ordering robust to a 2.39× reach-duration mismatch). The inter-array anterior-posterior offset (∼2 mm) was registered to the ICMS current-threshold trough, the electrophysiological landmark common to both implants, rather than to raw stereotaxic coordinates. The within-array somatotopy map is the primary registration (it carries the toward-3a projection and the sink/source colocalization, both computed within a single subject); the cross-subject stitch into one premotor-to-3a crescent (Fig. 5D,E) is a secondary, consistency-checked claim. The ∼2 mm is a centre-offset proxy for the true inter-focus distance, so the implied ∼16 mm/s is conservative. Electrode pitch was 400 µm; inter-subject cortical variation is an acknowledged physiological and surgical coregistration uncertainty.

##### Pre-spike control

To test whether an organized δ wave precedes the spike sweep, the PRE window was widened to at least one δ half-cycle and was not envelope-anchored. Four discriminators separate a translating wave from a static spatial offset: phase-gradient directionality (PGD, the planar coherence of the phase-flow field defined above) versus a spatial-shuffle null; the anterior-to-posterior front-translation lag (finite for a real wave, near zero for a static offset); alignment of the instantaneous flow direction to the anterior-posterior axis; and the divergence sign in PRE versus POST. The no-preceding-wave verdict held across anchor choice, PRE-window width, amplitude weighting, and ±50 ms detection-anchor jitter. The single calibrated point against a strict field-shadow reading is that the discriminating sink/source polarity is already present in the PRE window, before the spikes fire (TY 56% PRE-to-POST sign match, p = 0.016; JL 92%, p = 0.003). This early polarity reflects the orientation of a co-traveling field rather than a field-to-spike drive: the sink/source sign is established before the spike peak, yet the organized δ propagation still does not precede the spike sweep, so an early polarity does not imply that the field organizes or drives spiking.

##### Nulls and robustness

Shuffle nulls were used throughout; the quick-capture-versus-correction contrast additionally carried controls for movement speed-peak, correction isolation (inter-event interval > 300 ms), within-episode correction position, detection-anchor jitter (±50 ms), window width, amplitude weighting, and cross-subject clock comparability. Movement-detection jitter is controlled but not fully eliminated, so part of the reduced correction-wave coherence may reflect residual jitter rather than a pure movement-type difference.

### Statistical analysis

Unless noted otherwise the significance threshold was α = 0.05 and all analyses were performed in MATLAB. We report two layers of statistics that serve different purposes and should not be conflated. On-figure significance brackets are per-subject descriptive tests that treat events as independent, used to show that an effect is visible and replicates within each animal on the raw per-event distributions. Inferential cross-subject claims, where made, come from a linear mixed-effects model (‘fitlm’, restricted maximum likelihood). Cross-subject mixed-effects inference is applied to the per-event jPCA rotational metrics only, which is the one quantity in continuous per-event form with sufficient n (≈330 events across both subjects) to estimate the subject and episode variance components. Its episode random intercept is material (≈9% of residual variance), so omitting it would understate the standard errors. The tangling, PETH, and subspace analyses are evaluated per subject (with effect sizes), and their cross-subject conclusions rest on independent replication of the same ordering in both animals rather than on a pooled fit, because their respective units (per-timepoint Q within a condition; per-unit firing on two physically distinct arrays) are not exchangeable across subjects and would constitute pseudoreplication if pooled naively. The PETH population-pooled row is descriptive, not an inferential pooling across subjects.

Mixed-effects model (jPCA per-event metrics). The inferential claims for the jPCA per-event metrics are fixed-effect contrasts from a linear mixed-effects model fit in MATLAB (‘fitlm’) by restricted maximum likelihood (REML) to the per-event values pooled across both subjects. In Wilkinson notation the model is: Metric ∼ MovementType + (1 | Subject) + (1 | Episode), with movement type a fixed effect (reference level Quick-Capture) and random intercepts for subject and for episode; fixed-effect F-tests use the residual degrees of freedom (the ‘fitlmè default). REML is used rather than ordinary maximum likelihood because it estimates the random-effect variance components with approximately no bias. The random episode intercept models the non-independence of events drawn from the same pursuit episode (episode key = the subject, video-label, episode-in-video tuple). The pre-specified contrast is Correction versus Quick-Capture: frequency Co − QC = −0.506 Hz (omnibus F(4,329) = 26.4, p = 5×10⁻¹⁹; contrast F(1,329) = 28.2, p = 2×10⁻⁷); amplitude Co − QC = −0.452 a.u. (omnibus F(3,301) = 17.5, p = 2×10⁻¹⁰; contrast p = 1×10⁻⁹). Each contrast is cross-checked with an episode-block bootstrap (resample whole episodes with replacement, 2000 iterations, recomputing the mean difference): frequency 95% CI [+0.35, +0.69] Hz, amplitude 95% CI [+0.35, +0.65] a.u., both excluding zero and sign-concordant with the model. Because the bootstrap resamples already-computed per-event values rather than refitting jPCA, it is free of the bootstrap-distribution-variance-deflation problem that disqualifies bootstrapping the eigenvalue for inference. Vacillation is excluded from the cross-type omnibus, post-hoc, and mixed model and reported descriptively only; JL Vacillation (n = 9) additionally carries an explicit small-n caveat and is never a point claim.

Per-subject descriptive tests for the jPCA metrics (Fig. 3F,vi-vii). As a within-subject check that the ordering is visible on the raw per-event distributions, frequency and amplitude were also tested separately within each subject, treating events as independent: a Kruskal–Wallis omnibus with Dunn’s post-hoc (Bonferroni-corrected over the ten pairwise comparisons) for frequency, and a one-way ANOVA with Tukey–Kramer HSD for amplitude. A non-parametric omnibus is used for frequency because the per-event phase-rate distributions are right-skewed and heteroscedastic across types (notably the heavy-tailed Correction distribution, CV ≈ 0.7–1.0); the detrended amplitude distributions are approximately symmetric, so a parametric family is appropriate and Tukey–Kramer handles the unequal group sizes. Because these tests do not model within-episode non-independence and do not pool across subjects, significance is not marked on these two panels and the reported inference for them is the mixed-effects model below.

Tangling (Fig. 3E,ii-iii). Distributions of log₁₀(Q) for QC vs DP episodes (Fig. 3E,ii) were compared by two-sample Kolmogorov–Smirnov test, chosen because Q is positively skewed even on log scale and the KS statistic is sensitive to both location and shape differences. With sample sizes in the thousands of timepoints, asymptotic p values past 10⁻¹⁰ are not numerically meaningful; we therefore report a p < 10⁻¹⁰ bound alongside the KS D statistic and Cohen’s d on log₁₀(Q). Per-event median Q per movement type (Fig. 3E,iii) was compared by Wilcoxon rank-sum (Mann–Whitney U), with effect size r = Z/√(n₁ + n₂). The QC-Initial vs Correction contrast is the pre-specified primary comparison and is reported uncorrected; the two secondary contrasts (DP-Initial vs QC-Initial; Reset vs QC-Initial) are reported Bonferroni-corrected across that two-test family. We chose rank-sum over a t-test because per-event Q distributions are heavy-tailed within each movement-type cell, and over a KS test because the secondary question was specifically a location shift rather than a shape difference.

PETH heatmaps (Fig. 4). The inferential class-selective contrasts were used to test whether QC firing in each class’s dominant window is reduced relative to each of the three other movement-type columns. Test: paired Wilcoxon signed-rank on each unit’s absolute mean normalized firing rate in its dominant window (Pre / Extension / Post, classified by argmax of QC-column window means), comparing the QC column to each reference column (Correction, Reset, DP-initial). Class assignment is held fixed by QC across contrasts so per-class n is constant (TY 27 / 90 / 46; JL 18 / 79 / 36; signed-rank df = n − 1 per class). 95% CIs on the median paired difference (and on its percent-reduction expression) were estimated by 2000-iteration paired-median bootstrap (resample units with replacement; seed fixed for reproducibility). All p-values reported uncorrected; the graded hierarchy described in Results is supported by the population-pooled row in both subjects (all six pooled p < 10⁻³).

Reporting conventions and uncertainty. Box plots show the median and interquartile range with whiskers at 1.5× IQR; points are per-event values; error bars and 95% confidence intervals were obtained by the bootstrap procedures stated above. Non-parametric tests were chosen wherever per-event distributions are right-skewed and heteroscedastic (assessed by inspection of the distributions rather than by a formal normality test); parametric tests (one-way ANOVA / Tukey–Kramer) are used only for the approximately symmetric detrended-amplitude distributions. Multiple-comparison corrections are as stated per test: Bonferroni over the ten pairwise jPCA frequency post-hocs and over the tangling two-test secondary family; the pre-specified primary contrasts (jPCA Correction-vs-QC; tangling QC-Initial-vs-Correction) and the PETH class contrasts are reported uncorrected.

Per-class table in Supplementary Table S1.

Population pooled; each unit contributes its own dominant window.

| Subject | QC – Correction | QC – Reset | QC – DP-initial |
| --- | --- | --- | --- |
| TY (n=163) | +15% [+11%, +17%]<sup>***</sup> | +9% [+7%, +11%]<sup>***</sup> | +2% [-0%, +5%]<sup>***</sup> |
| JL (n=133) | +20% [+16%, +24%]<sup>***</sup> | +16% [+12%, +20%]<sup>***</sup> | +5% [+3%, +9%]<sup>***</sup> |

Values are median percent reduction from QC (95% bootstrap CI). Significance: \* $p < 0.05$ , \*\* $p < 0.01$ , \*\*\* $p < 0.001$ ; all pooled cells $p < 0.001$ (\*\*\*).

Pursuit-band partition counts. The Correction-anchored peri-event partition (Methods, Correction-anchored peri-event partition) classifies each unit as engaged / disengaged / off-band using a deterministic two-criterion rule on smoothed Correction and QC-Initial PETH traces. The partition is descriptive (no inferential test); we report the per-subject counts as the summary, together with the row span occupied by the engaged class under a quick-capture-latency ordering.

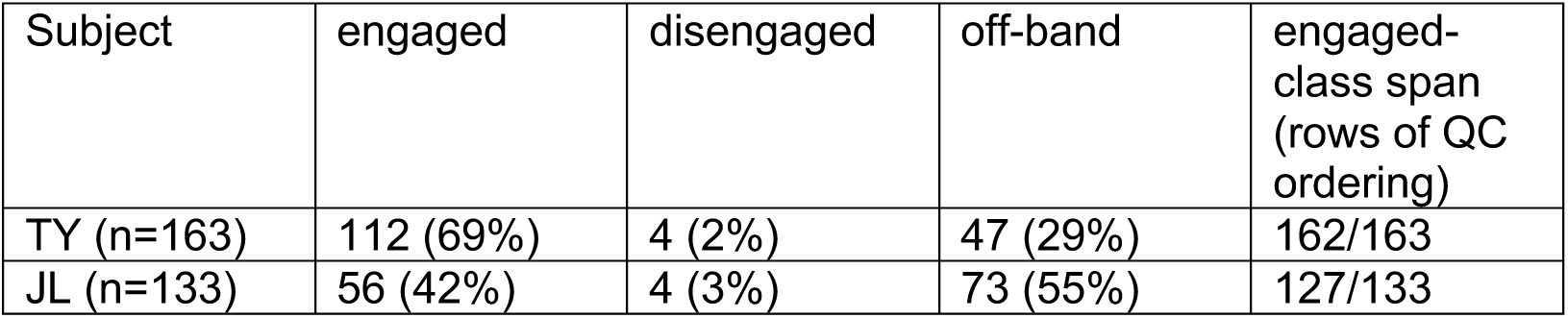

The span is the row-index distance between the first and last engaged unit under the quick-capture-latency ordering. In both subjects the engaged class spans nearly the whole population (162 of 163 rows in TY; 127 of 133 in JL), indicating interleaving with off-band rows rather than a contiguous block. This is the operational expression of the interleaved-vs-block geometry described in Results.

### Kinematic-confound controls

i. Subspace / PLI. Because within-reach corrections are kinematically smaller than initial reaches (shorter, slower, executed in flight), we further verified that the near-zero correction lead reflects a genuine difference in preparatory engagement rather than a movement-magnitude artifact. As the PLI is read from event-averaged occupancy and has no per-event value, we used each event’s peak preparatory-subspace occupancy magnitude in the pre-detection window (−0.75 to 0 s) as a per-event surrogate, the quantity such a confound would suppress, since the magnitude of preparatory re-engagement scales with the size of the required correction (Ames et al. 2019). Regressing the log preparatory-occupancy magnitude on per-event reach kinematics (reach extent, peak speed, and path length) together with movement type, the deficit in preparatory occupancy for corrections relative to initial reaches remained significant after partialling out the kinematic covariates in both subjects (TY p = 0.003; JL p = 1.5 × 10⁻⁴; movement-type partial F-test p = 0.005 and 3 × 10⁻⁷; reproduced by a gamma-family generalized linear model). The reduced preparatory engagement of corrections is therefore not a movement-magnitude artifact. This magnitude surrogate distinguishes initial reaches from non-initiations but not resets from corrections, consistent with magnitude and timing (PLI) indexing complementary facets of preparatory re-engagement. A complementary matched-kinematics analysis, subsampling corrections and initiations to overlapping distributions of reach extent and peak speed and recomputing the occupancy lead, corroborated the ordering in TY but could not be applied in JL, whose corrections and initiations are kinematically near-disjoint (mean reach extent 2.8 versus 7.0 cm) and so lack the common support a matched comparison requires.
ii. jPCA. Because within-pursuit Corrections are mechanically the shorter, slower, mid-flight follow-up movements (median reach extent ≈ 2.7–2.9 cm versus ≈ 7.0 cm for quick-capture initiations in both subjects), we verified that the slower, smaller-loop Correction signature is not a kinematic-magnitude artifact. For each subject we regressed the per-event frequency (raw signed Hz) and log rotational amplitude on movement type together with per-event reach kinematics (reach extent, peak hand velocity, and total path length), with Quick-Capture as the reference level, across a conservative family of five model specifications (movement type alone, each single covariate added, and all three). In JL the Correction-versus-Quick-Capture contrast survived full kinematic control for both metrics (frequency Co − QC = −0.498 Hz, p = 0.011; log amplitude = −0.199, p = 0.0011) and reach extent did not significantly predict JL frequency (p = 0.55). Class and kinematics contribute largely orthogonal variance, so the Correction effect is independent of movement extent. In TY the pairwise Correction contrast was collinear with reach extent and absorbed once it was included (frequency full-model β = −0.101, p = 0.48; log amplitude = −0.026, p = 0.66), so in TY the slower-and-smaller signature cannot be statistically separated from the smaller mechanics of the movement. A complementary matched-kinematics analysis was not applicable in either subject: Corrections and quick-capture initiations are kinematically near-disjoint, leaving only 9 of 88 (TY) and 4 of 58 (JL) pairs within a 0.5-SD caliper, below the n ≥ 10 threshold. This disjointness is itself informative. Corrections occupy a distinct, smaller-extent kinematic regime, so the altered correction dynamics cannot reflect overlapping kinematic variation. However, it also bounds the regression: with little common support, the models adjust for kinematics partly by extrapolating between the two distributions. We therefore read the surviving JL contrast as an existence proof of kinematics-independence rather than as a claim that the relation is linear throughout.
iii. Tangling. To verify that the movement-type-graded Q signature is not a trivial consequence of kinematic differences across movement types, we fit per-subject linear models, in Wilkinson notation log₁₀(Q) ∼ movement_type + peak_velocity + path_length + reach_extent, with QC-Initial as the categorical reference level; the movement-type coefficient reported in Results is therefore the adjusted difference in log₁₀(Q) between corrections and quick-capture initial reaches, holding the kinematic covariates fixed. Per-event kinematic scalars were extracted from the 3-D hand position and velocity timeseries over each event’s [−250, +150] ms window: peak hand velocity (90th-percentile speed, robust to single-frame tracking spikes), total path length, and 3-D reach extent from the position at detection. Models were fit on n = 157 events (TY) and n = 108 events (JL) after excluding episodes in which the video started after reach onset, events with NaN velocity, and a TY failed capture episode in which tangling values exceeded a 3xIQR criterion. To guard against covariate collinearity (the three kinematic scalars are pairwise correlated r = 0.77–0.90 within the QC + Correction subset), we additionally fit each kinematic covariate alone in turn, plus a no-covariate baseline; the kinematic-confound disposition is determined jointly across this five-model family.

### Intracranial microstimulation (ICMS)

ICMS was performed in calmly resting, unconstrained marmosets prior to falling asleep. Current (biphasic pulses, 3–60 µA range with 60 µA maximum observed; 500 µs pulse width, 55 µs interphase interval, 200 Hz, 100 ms burst) was delivered to a single randomly chosen electrode at a time through the Nano-Stim headstage, coordinated by the Trellis software suite and the grapevine neural interface box (Ripple Neuro). Stimulation parameters were chosen to be in line with those used in our lab previously and with those found in the literature (Burman et al. 2008; Burish et al. 2008; Huffman 2001), however we were largely able to use much smaller threshold currents to evoke effects in the muscles due to the marmosets being calmly resting rather than sedated as in standard approaches. The procedure generally took place in 2-3 hr sessions over multiple nights. If proceeding smoothly, such a session would allow time to sample ∼32 electrodes. For each electrode we would begin at 15 µA and the experimenter would use fingertips to gently probe the marmoset’s skin contralateral to the array to identify the muscle that was responsive to the stimulation. Once a muscle or body part was identified current was reduced until the threshold at which current would no longer produce muscle activation. If no muscle was found to respond at 15 µA, current would be increased in small increments to identify a responsive muscle and the process of reducing to a threshold was repeated. If no responsive muscle was identified by 60-100uA, the electrode was labeled as unresponsive. Notes from these sessions were then used to reconstruct the somatotopic maps for each array (Fig. 5A).

### Cortical area designation

The ICMS results were then used to identify the most likely placement of the arrays and each electrode’s position within known electrophysiological and anatomical features of marmoset sensorimotor cortex (Burman et al. 2008; Burish et al. 2008; Huffman 2001). Both arrays were implanted to target forelimb area of primary motor cortex in marmoset at (9.8AP,4.5ML) as estimated by a combination of (Paxinos et al. 2012) and (Burman et al. 2014). After implantation the primary electrophysiological feature that allowed for estimation of placement within sensorimotor cortex was minimum stimulation threshold as described above, and averaged across medial lateral spanning columns of electrodes in both marmosets. In addition to percentage unresponsive electrodes in those same medial lateral spanning columns, we found a clear pattern of low thresholds that were flanked anteriorly and posteriorly with less excitable cortex. The width of the electrophysiological trough in thresholds was consistent with the width of M1/3a as reported in previous literature (Burman et al. 2008; Burish et al. 2008; Huffman 2001). In TY, posterior to this trough we found units that expressed responses to tactile stimulation consistent with the most posterior TY electrode placement in primary sensory area 3b (Krubitzer and Kaas 1990). In JL, anterior to this trough we found a cluster of unresponsive electrodes and electrodes with high thresholds, consistent with anterior most JL electrodes placement in dorsal premotor cortex (PMd).

## Use of AI tools

Claude and Gemini were used in preparing code and in drafting prose from initial versions written by JW. All authors take responsibility for the results reported. The marmoset illustrations in Figures 1A and 5A were produced with NanoBanana by iterating on initial sketches by JW with style references.

## Acknowledgements

We would like to thank Carmen Eggleston and other members of the Hatsopoulos lab for help with experiments and marmoset care. We would also like to thank the University of Chicago veterinary staff for exceptional marmoset care and surgical assistance.

## Supplementary Information

Supplementary Video S1. Example prey-capture episodes with simultaneous population dynamics. Six example episodes from marmoset JL: three quick captures, then three dynamic-pursuit episodes. Each clip is the animated form of the panels in Fig. 2A and Fig. 3A: synchronized behavioral video, the neural population trajectory in the low-dimensional state space with a moving dot marking the current time point, hand-position traces, and the simultaneously recorded spike raster, with a cursor sweeping the time axes.

Supplementary Video S2. Single-trial spike–δ overlay near peak hand extension. Single-trial overlays of the δ-band (1–4 Hz) LFP field (greyscale squares, one per electrode, dark = trough) and concurrent firing rate (circle brightness, white = high) across peak hand extension. The video plays two example events in sequence: first marmoset TY (episode 80), then marmoset JL (video 17, episode 1). Each panel’s lower plot shows concurrent hand kinematics with a cursor marking the displayed frame; playback is slowed (TY 5.0×, JL 4.5×). Every square carries a measured δ-LFP value from its own electrode; no values are interpolated or spatially smoothed. These are the animated versions of the static panels in Fig. 5D,i-ii.

**Table S1.**
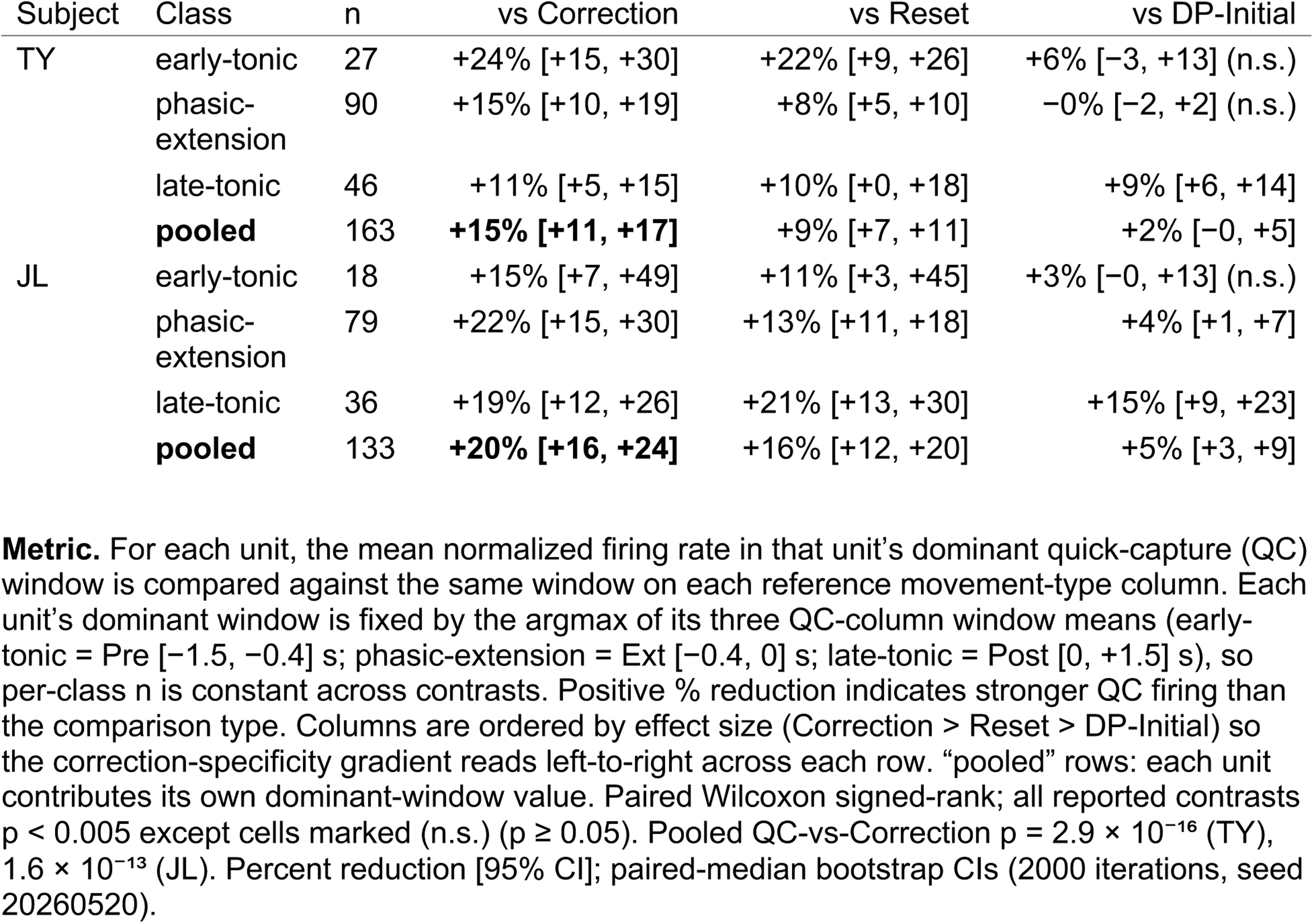
Class-selective firing-rate reduction (quick capture vs other movement types)

**Fig. S1.**
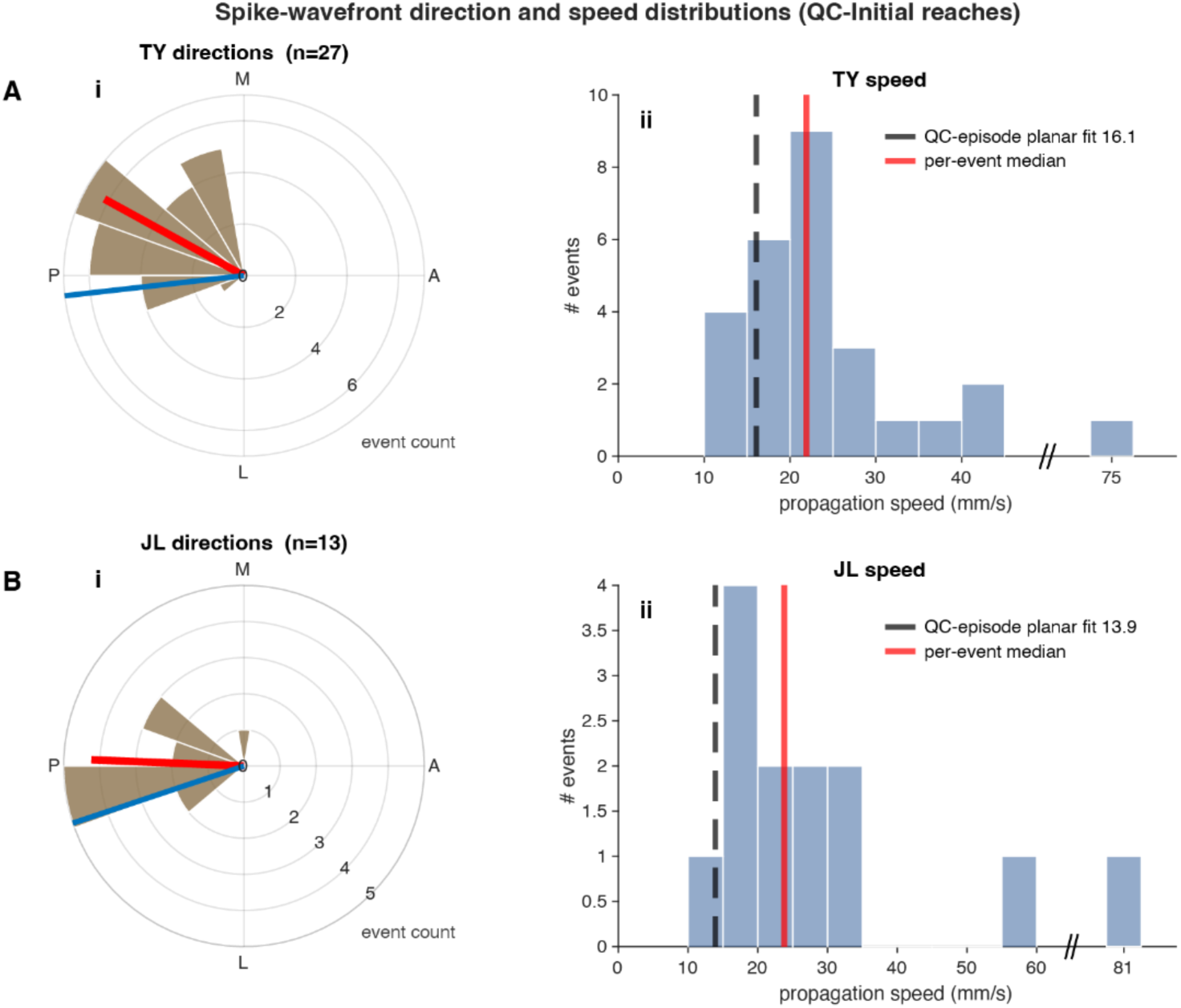
Spike-wavefront propagation direction and speed distributions for quick capture reaches Per-event distributions for QC-Initial reaches, shown for both subjects (A, TY, n = 27 events; B, JL, n = 13). (i) Direction roses. Angular histograms of single-event spike-sweep direction, 18 bins of 20°. Wedge length is the number of events in that direction bin. The red radial line is the circular resultant vector: its direction is the mean sweep direction and its length is the resultant magnitude R (0 = uniformly distributed, 1 = all events identical), plotted as a fraction of the radial axis. The blue radial line marks the array-centre→area-3a axis for reference. Axes are the cardinal anatomical frame (A, anterior; M, medial; P, posterior; L, lateral); JL angles are rotated by 135° into this common frame so the two subjects are directly comparable. (ii) Speed histograms. Per-event propagation speed, 5 mm/s bins. The x-axis is broken (‘//’) to accommodate a single far outlier per subject (TY 75, JL 81 mm/s); the outlying event is plotted beyond the break at its own labelled tick, so every event is shown and counted and no data are clipped. As marked in the legend, the red line is the per-event median (TY 21.9, JL 23.8 mm/s) and the black dashed line the median per-episode planar-fit speed over significant quick-capture episodes (TY 16.1, 19 of 28; JL 13.9, 9 of 13 mm/s), as quoted in the main text and annotated in Fig. 5B. Note the change of unit: the histogram is per event, the dashed line per episode. Per-event speeds are inflated where the planar fit is poor (Spearman ρ between per-event R² and speed = −0.91 TY, −0.94 JL), so the dashed line is a non-attenuated reference, not a summary of the plotted data; restricting to individually significant events, a separate event-level test, gives 19.9 mm/s in TY (n = 19) and 15.8 in JL (n = 5). Plotting the full distribution leaves JL’s reference line below all but one of the plotted events, the expected direction of this bias at n = 13.

